# A heart releasing neuropeptide that synchronizes brain-heart regulation during courtship behavior

**DOI:** 10.1101/2025.03.31.646145

**Authors:** Hongyu Li, Shenjie Cheng, Yijun Niu, Tongxin Diao, Qiruo Zhang, Wei Zhang

**Affiliations:** State Key Laboratory of Membrane Biology, School of Life Sciences, IDG/McGovern Institute for Brain Research, Tsinghua University, Beijing 100084, China; Tsinghua-Peking Center for Life Sciences, Beijing 100084, China

## Abstract

Distinct internal states drive varied animal behaviors, yet the mechanisms by which non-neuronal factors encode these states remain largely unknown. Here, we show that cardiac activity regulates internal mating states through a conserved brain–heart axis in male flies and mice. In Drosophila, a ppk23-P1 pathway triggers heart rate acceleration upon female perception via crustacean cardioactive peptide, while the heart secretes ion transport peptide that feeds back onto P1 neurons to enhance courtship. Ejaculation rapidly decreases heart rate via Corazonin, mitigating prolonged tachycardia. In mice, a conserved Substance P-TacR1 pathway modulates courtship. Finally, we developed a computational framework to decode internal mating states from cardiac physiology, revealing distinct cardiac signatures. Our findings unveil a novel role for heart-derived neuropeptides in internal state regulation, elucidate a positive feedback loop between the heart and brain, and demonstrate the evolutionary conservation of the brain-heart axis in orchestrating dynamic behavioral and physiological states.

## Background

Central nervous system is immersed in dynamic and multidisciplinary processing demands, shaping the ongoing organization between external stimuli and behavioral responses, herein referred to as’internal states’^1–3^. Internal states play a pivotal role in driving behavioral decisions, particularly in fundamental processes such as feeding, mating, and sleep. These states are considered to be encoded by dynamic neural activity, but the broader interaction between multiple organ systems remains poorly understood. Recent studies have revealed that feedback from certain organs plays crucial roles in a variety of behaviors^4–6^. For examples, gut sensing have been reported to regulate feeding^7^, immune response^8–10^, and reproduction^11–15^. While blood pressure sensing and related sensory mechanisms have been recently discovered^16,17^, the cardiac system remains less investigated in terms of its impact on animal behavior and internal states.

The heart is an enduring symbol of intense feelings across many cultures. Cardiac activity is highly responsive to and shaped by different internal states across species^18,19^. Human exert distinct patterns of cardiac activity during a variety of emotional states^20–23.Cumulative^ evidences outline that the brain-heart connection highly affects human health from genetic and structural perspectives ^24–27^. Beyond its role in circulation, the heart is increasingly recognized as an active signaling organ, capable of releasing various bioactive molecules that influence neural activity and behavior^28–30^. Among these molecules, neuropeptides are widely expressed in the heart^27,31^, yet their roles in modulating neural processes and behavioral states remain largely unexplored. Notably, emerging studies revealed the causal link between the brain and the heart in different emotional and physiological states, for instance, the optically induced tachycardia elicited anxiety-like behaviours in a context dependent manner^32^, suggesting that cardiac activity is sufficient to induce changes of internal states. Growing evidence supported that cardiac system could interact with central nervous system by diverse regulatory pathway^17,30,33^, yet the encoding mechanism of internal states by such a brain-heart axis is still elusive. Despite growing recognition of the heart’s capacity to communicate with the brain and other organ, how these cardiac-derived signals contribute to the regulation of internal states is still an open question. Sexual activity, one of the most intense and conserved behaviors, has long been associated with cardiac activity. Based on the prominent perception “butterflies in the stomach”, studies have revealed the strong correlation between cardiac activity and sexual arousal in both sexes^26^. For instance, strangers with higher heart rate synchrony during dating predicts more attraction without consciously noticing^34^. In animals, males often exert higher heart rate and cardiac function during courtship across taxa^35–38^. While the central nervous system’s role in regulating diverse mating states in male animals has been extensively studied, the involvement of the heart in courtship and evolution, the molecular and neural basis of brain-heart cross-talk remain unclear. It is still uncertain whether, and how, the heart contributes directly to the regulation of internal mating states.

To bridge these gaps, we investigate how cardiac activity is linked to internal mating states and the role of the brain-heart axis in regulating sexual behavior. We show that distinct cardiac activity patterns emerge in response to different mating states, with neuropeptides released from cardiomyocytes playing a critical role in modulating male competitiveness during courtship. These findings suggest that the heart is actively involved in regulating courtship behaviors through its signaling molecules. Additionally, we highlight the evolutionary significance of these processes, as courtship behaviors are crucial for reproductive success and provide a competitive advantage. Using mice and fruit flies as model organisms, we explore the conserved mechanisms of heart-derived signals across species, shedding light on the role of the brain-heart axis in driving reproductive success and individual fitness.

## Results

### Female perception flips internal mating states to trigger cardiac activation in male flies and mice

We first examined how male flies respond to female cues under different internal mating states. Using a tethered preparation (Fig. 1A), we recorded the heart rate of male flies and found that flies in distinct mating states exhibit different frequency distributions of cardiac activity—successful mating is associated with a lower dominant frequency, whereas failed mating corresponds to a higher frequency (Fig. 1B). When we presented various external stimuli, only contact with a female fly elicited a transient acceleration of heart rate; stimulation with a metal probe or a male fly had no such effect (Fig. 1C, D, Extended Data Figure 1A). Notably, naïve males exhibited a significant heart rate increase upon female exposure, while mated males did not, indicating that the cardiac response reflects the male’s internal mating state.

**Figure 1.**
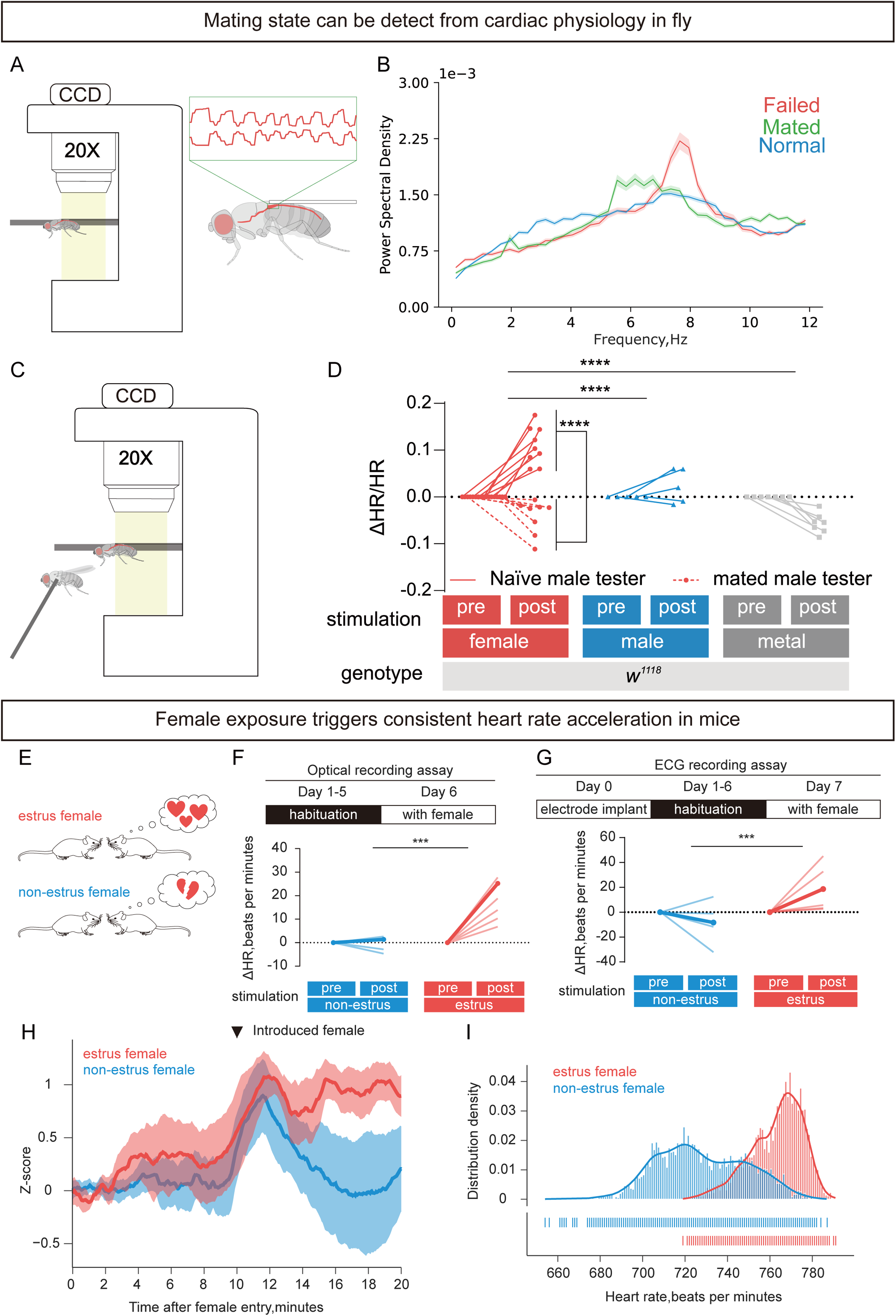
Female perception flips internal mating states to trigger cardiac activation. **A**, Schematic illustration of the heartbeat recording. Flies are tethered with optical adhesive on a coverslip. The heart chambers and aorta are highlighted with red color. **B**, Cardiac activity from distinct mating states exhibit different distribution pattern in frequency domain. Here the mating states separate the main frequency to lower (success mating) or higher (failed mating) range. **C**, Schematic illustration of the heartbeat recording system with different stimulations (male, female, or metal probe). **D**, Flies under distinct mating states respond differently to female cues: naïve males exhibit a significant heart rate acceleration upon female exposure, while mated males do not. The y-axis represents the relative heart rate change (ΔHR/HR). Two-way ANOVA followed by Tukey’s test, p<0.0001. Female perception accelerates the heart rate of a male fly. Compared with a metal probe or male flies, touch on female flies can induce heart rate acceleration. Two-way ANOVA followed by Tukey’s test, p<0.0001. **E**, Schematic paradigm for recording heart rate in mice during social interactions with either estrus or non-estrus females. **F**, The heart rate change of a male mouse is significant higher after coupling with an estrus female mouse than with a non-estrus female. Heart rate is measured by the optical recording assay. Two-way RM ANOVA followed by Šídák’s multiple comparisons test, p = 0.0003. **G**, Heart rate change of a male mouse is significantly higher after coupling with an estrus female mouse than with a non-estrus female. Heart rate is measured by the ECG recording assay. Two-way RM ANOVA followed by Šídák’s multiple comparisons test, p =0.0051. **H**, Average heart rate (z-score) of male mice before and after coupling with a female mouse (estrus or non-estrus) during whole social interaction session. **I**, The heart rate distribution in male mice after coupling with an estrus female is significantly shifted higher compared to coupling with a non-estrus female (unpaired t-test, p < 0.0001).

A similar phenomenon was observed in male mice. We utilized both optical recording and electrical recording assays, which are widely employed methods for heart rate measurement^18,32^ (Figure 1 E, Extended Data Figure 2 A,B), to monitor the cardiac activity of male mice during sexual behavior. We first ensured the stability of the ECG signal after electrode implantation for 5 days (Extended Data Figure 2 C, D) and verified that the mice maintained normal body weight post-surgery (Extended Data Figure 2 E, F). In the female exposure assay, we recorded the heart rate of male mice using either the optical method or ECG recording for 10 minutes in their home cage to establish a baseline for the heart rate in resting state. Subsequently, we introduced female mice in either estrus (red) or non-estrus (blue) into the mating chamber. Male mice interacting with estrus females exhibited robust and prolonged heart rate acceleration, whereas non-estrus females induced only a transient increase in the heart rate (Figure 1F-I). This change in the cardiac activity aligns with the behavioral outcomes: male mice typically engage in active exploration of unfamiliar female mice, and initiate courtship behavior even without the response from the female. However, the later steps of the mating behavior are elicited only by the arousal states triggered from estrus females, whereas non-estrus females inhibit further courtship attempt from male mice by rejecting them.

These findings underscore the involvement of cardiac activity in modulating male mating behavior and reflecting internal mating states. Building on this, we next examined how neuronal dynamics interface with the cardiac system, exploring molecular pathways and circuit mechanisms that drive these interactions.

### A ppk23-P1-CCAP pathway mediates female induced cardiac response in fruit fly

To identify the neural basis of the female-induced cardiac response in male flies, we investigated the role of ppk23^+^ neurons, which are known to transduce female chemical profiles when a male touches a female with their foreleg tarsi^39^. In *ppk23* mutant flies, exposure to a female did not elicit heart rate elevation (Figure 2A). The ppk23^+^ neurons on the tarsi comprise two types: F (female-responding) and M (male-responding) cells^40^. By selectively expressing the inwardly rectifying potassium channel Kir2.1 in F cells, we silenced these female-responding cells and observed that female contact no longer accelerated the male heart rate (Figure 2B), indicating that female pheromonal cues trigger heart rate acceleration through ppk23^+^ sensory neurons.

**Figure 2.**
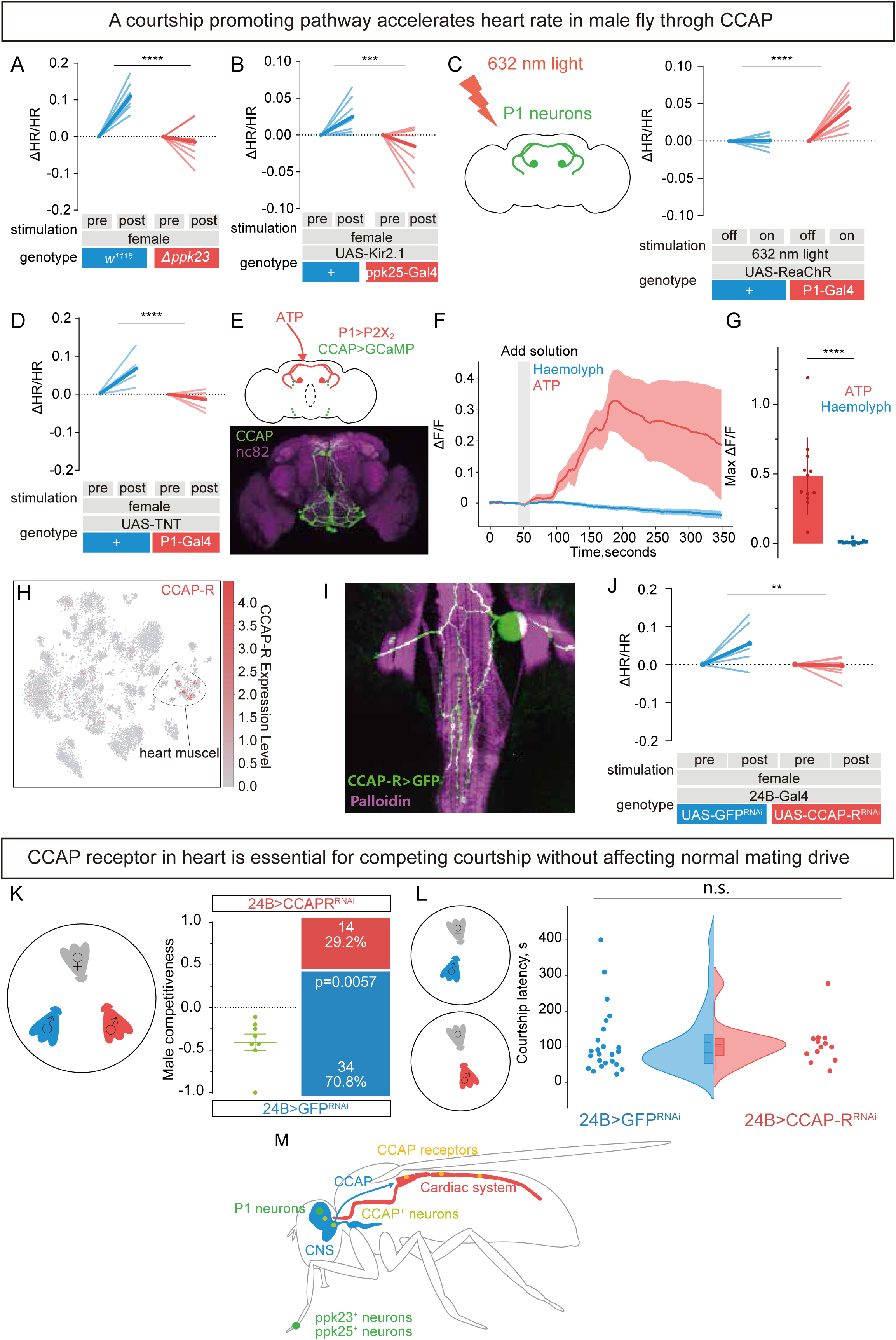
Heart rate acceleration is mediated through a courtship-promoting circuit that enhances male competitiveness in flies. **A**, Mutation of the pheromone-sensing receptor gene (*ppk23*) abolishes the cardiac response to female stimulation (two-way ANOVA with Tukey’s test, p < 0.0001). **B**, Silencing F cells (*ppk25*^+^cells) blocks the cardiac response to female stimulation. Two-way ANOVA followed by Tukey’s test, p=0.0005. **C**, The heart rate of male flies increases in response to 632nm light. Two-way analysis of variance (ANOVA) followed by Tukey’s test, p<0.0001. **D**, Blockage of the synaptic transmission of P1 neurons inhibits the female induced heart rate acceleration. Two-way analysis of variance (ANOVA) followed by Tukey’s test, p<0.0001. **E-G**, Activation of P1 neurons by adding ATP induces calcium influx in CCAP neurons. Unpaired t-test, p<0.0001. **H**, The scRNA sequencing data revealed that CCAP receptors are broadly expressed in heart tissue and is enriched in cardiac muscle cells, data from fly cell atlas (https://www.flycellatlas.org/). **I**, Representative immunostaining confirms the expression pattern of CCAP receptors in the male fly heart tube. **J**, Knocking down CCAP receptor (CCAP-R) on the heart muscle blocks the female induced heart rate acceleration. Two-way ANOVA followed by Šídák’s multiple comparisons test, p=0.0012. **K**, Knocking down CCAP-R on the heart muscle impairs the courtship competitiveness of male flies. Binomial test, p =0.0057. **L**, Knocking down CCAP-R on the heart muscle does not affects courtship latency of male flies, indicating the normal mating drive of male flies remains. Unpaired t-test, p = 0.8804. **M**, Illustrative figure of the brain-heart innervation in male flies that relays female pheromones. A ppk23/ppk25-P1-CCAP pathway that accelerates heart rate in male flies during courtship behavior.

**Figure 3.**
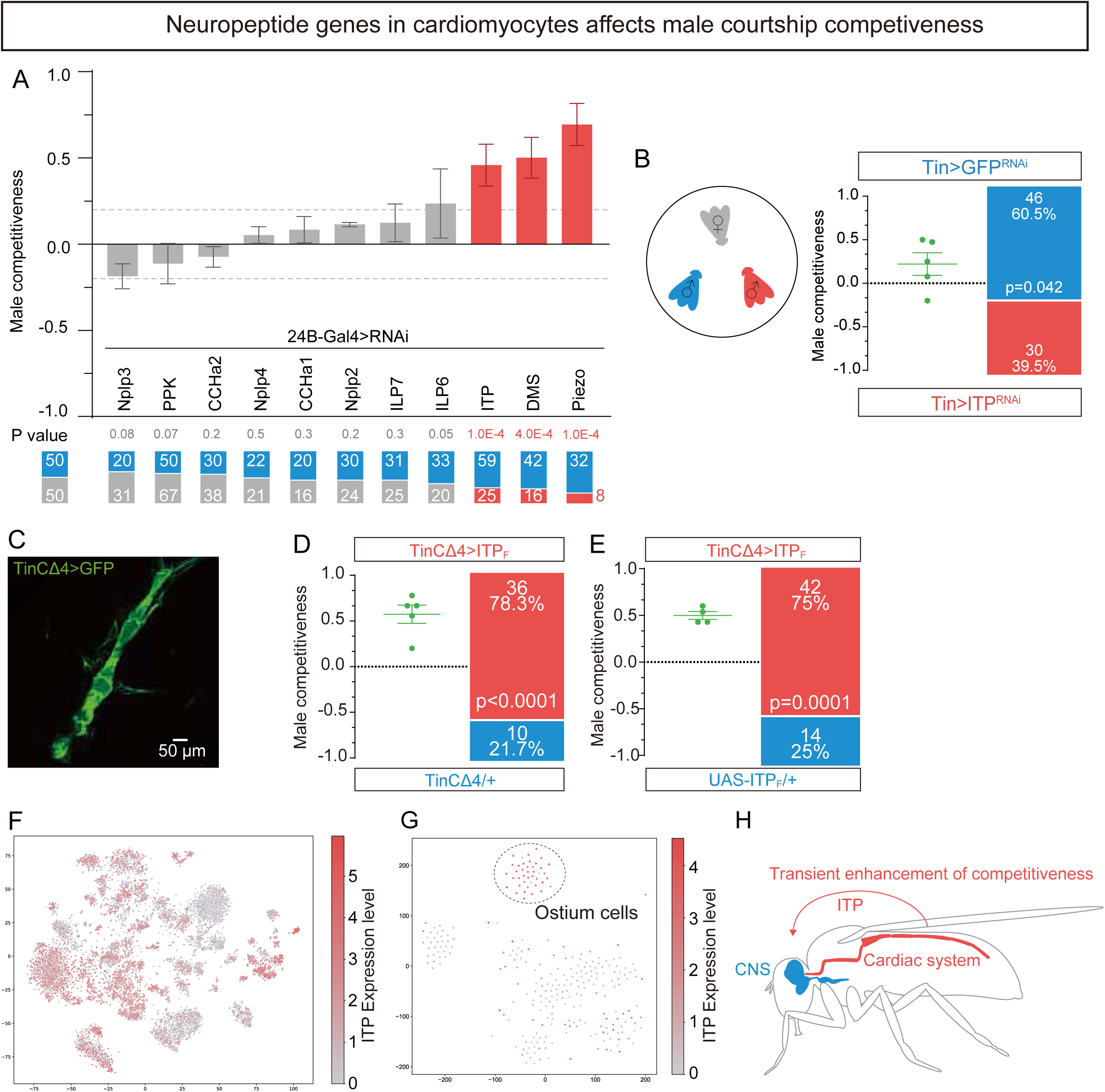
Neuropeptides produced by cardiomyocytes are essential for enhancing male competitiveness during courtship. **A**, A genetic screen of neuropeptide genes expressed in the heart identifies candidates that affect male mating competitiveness (binomial test; p values indicated in the figure). **B**, Knocking down *ITP* with an alternative driver confirms its essential role in promoting male competitiveness during courtship. Binomial test, p values are noted in the figure. **C**, Representative imaging shows the expression of the cardiomyocyte driver TinCΔ4-Gal4 in male flies. Scale bar is noted in figure. **D-E**, Overexpression of *ITP* gene in the cardiomyocytes enhances the male competitiveness. Binomial test, p values are noted in the figure. **F**, The scRNA sequencing data demonstrated that *ITP* is widely expressed in heart tissue and is enriched in several heart cell types, data from fly cell atlas (https://www.flycellatlas.org/). **G**, *ITP* is highly expressed specifically in Ostium cells and some heart muscle cells, data from fly cell atlas (https://www.flycellatlas.org/). **H**, During courtship scenario, *ITP* expression in cardiomyocytes and differentiated ostium cells is critical for male competitiveness.

We next addressed whether the response is relayed via the central nervous system. Decapitated flies did not exhibit heart rate changes upon female contact (Extended Data Figure 1B–C), indicating that the brain is essential for translating pheromone perception into a cardiac response.

P1 interneurons are reported to integrated external stimulation to initiate courtship behavior, we thus hypothesized that courtship-promoting P1 neurons in the brain mediate this response^41,42^. Using genetically encoded red-shifted channelrhodopsin (ReaChR), we selectively activated P1 neurons, and observed a marked increase in heart rate (Figure 2C). Conversely, blocking synaptic transmission in P1 neurons with tetanus toxin (TNT) abolished the female-induced cardiac acceleration (Figure 2D), highlighting the necessity of P1 neuron activation for this response.

Finally, we explored the downstream pathway by which P1 neurons relay this activation to the heart. Previous studies showed that Crustacean Cardioactive Peptide (CCAP) induces strong heart contractions in dissected fly hearts^43^. We thus tested whether P1 neurons activate the heartbeat via CCAP. In a dissected brain, we activated P1 interneurons by expressing P2X_2_ and adding ATP *ex vivo*, and observed that the Ca^2+^ level of CCAP^+^ neurons increased, while adding hemolymph alone could not induce such a response (Figure 2E-G and Extended Data Figure 3A, B). CCAP, upon release, binds to CCAP receptors on heart muscles, accelerating the heartbeat^43^. Consistent with these findings, single-cell RNA sequencing and immunostaining showed that CCAP receptors are broadly expressed in the heart, particularly in cardiac muscle cells (Fig. 2H–I). Finally, cardiac-specific knockdown of the CCAP receptor (CCAP-R) blocked the female-induced heart rate acceleration and impaired male courtship competitiveness without affecting the baseline mating drive (Fig. 2J–L).

Courtship and mating are challenging tasks that require multisensory integration and dedicated motor control ^42^. Besides, male flies continuously face competition from other males toward the same female in nature. We first tested whether the circulation system is essential for the courtship behaviors in male flies. The mutation of heartless (*htl*) leads to abnormal differentiation and migration of cardioblasts ^44^, resulting in defects of heart tube formation (Extend Data Figure 4A). The *hlt* males showed severe dysfunction in their courtship performance (Extend Data Figure 4B, C). To further investigate the relationship between cardiac function and courtship behavior, we knocked down TER94 with Hand-Gal4 to eliminate the whole heart tube (Extend Data Figure 4D). Notably, the absence of heart tube strongly dampened the courtship performance in male flies (Extend Data Figure 4E, F), confirming the essential role of the heart tube in courtship behavior. In contrast, eliminating the pericardial cells using RNA interference for the gene *klf15* had little impact on the courtship performance of the male flies (Extend Figure 4G-I).

**Figure 4.**
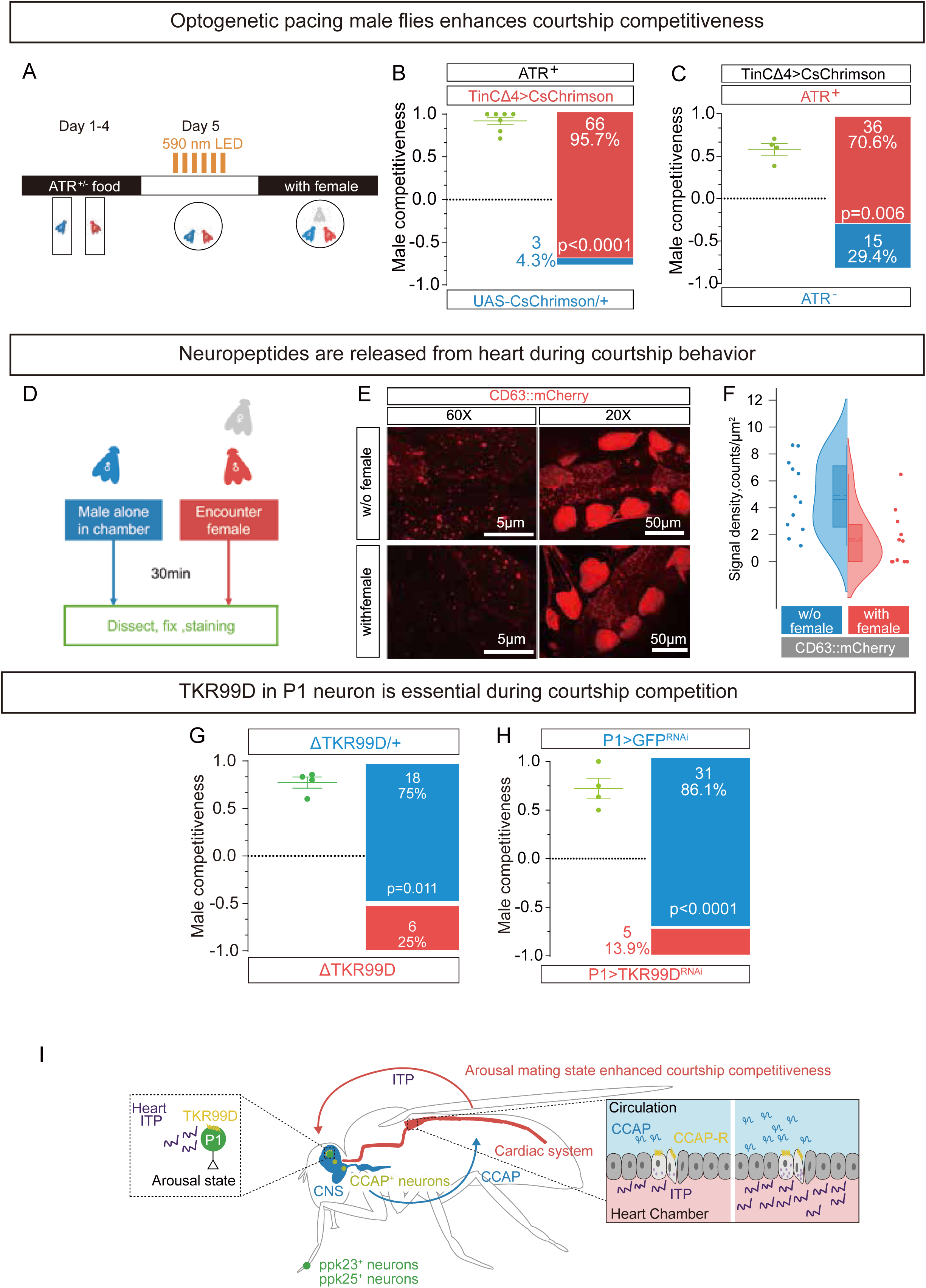
Neuropeptides released from cardiomyocytes enhanced male competitiveness by binding to TKR99D in courtship related neurons. **A**, Schematic illustration of the optical pacing assay. Flies were fed with all-trans-retinal (ATR) for 4 days after eclosion. At day 5, two males were introduced to the mating chamber and received pulsed light. After photoactivation, a female fly was introduced to these two male flies. **B-C**, Pulsed light activates the cardiomyocytes to enhance male competitiveness. Binomial test, p values are noted on the figure. **D**, Flies were placed in a courtship chamber with either a female or no stimulus for 30 minutes to allow successful mating. Subsequently, both mated and control flies were dissected, and their heart tubes were processed for exosome marker staining. **E**, Representative imaging of CD63 labeled signal in cardiomyocytes, scale bars are noted in figures. **F**, Quantification of CD63 signal indicates that female exposure induces vesicle release via exosomes in cardiomyocytes. Unpaired t-test, p =0.0323. **G**, Mutation of TKR99D impairs the competitiveness in male fly. Binomial test, p values are noted on the figure. **H**, Knocking down TKR99D in P1 neurons impairs male mating competitiveness. Binomial test, p values are noted on the figure. **I**, Schematic model of the feed-forward loop: female perception activates CCAP release into circulation, leading to an elevated heart rate that triggers ITP release from cardiomyocytes; the released peptides then bind to TKR99D in P1 neurons to sustain arousal states and enhance male competitiveness until successful mating.

As male flies with CCAP receptor (CCAP-R) knockdown in the heart could initiate courtship behaviors relatively normally (Figure 2L), this suggests that their basic mating drive and motivational states remain intact. However, in natural environments, animals often face intense competition for mates, where subtle differences in physiological and behavioral efficiency can determine reproductive success. To test whether cardiac CCAP signaling plays a role under such competitive conditions, we allowed these males to compete with control flies. Strikingly, males lacking CCAP receptors in the heart rarely succeeded against their competitors (Figure 2K). Moreover, they exhibited shorter mating duration (Extended Data Figure 3D), further confirming the critical role of cardiac responses in shaping courtship behavior, including copulation.

In summary, our results reveal a pathway in which P1 neurons act upstream of CCAP+ neurons, releasing CCAP that binds to heart muscle receptors, thereby driving female-induced heartbeat acceleration (Figure 2M).

### Cardiomyocyte-derived neuropeptides enhance male competitiveness during courtship

Having found the critical role of the heart in sexual behavior, we further wondered how the heartbeat acceleration promoted courtship performance. Considering the essential roles of the endocrine system that mediates the brain-organ communications in distinct internal states ^1,2^, we explored the recently released single cell RNA sequencing database^45^ to identify the neuropeptide genes that are preferentially expressed in the heart tissue, and selected several highly expressed neuropeptide genes as potential candidates (Extended Data Figure 5A-I). Next, we knocked down these genes in the heart with 24B-GAL4 and tested the males’ mating competitiveness. Knocking down *Dms* (Drosophila Myosuppressin, CG6440) or *Itp* (Ion transport peptide, CG13586) genes in the heart largely dampened the courtship competitiveness (Figure 3A). Additionally, the mechano-transduction channel gene *Piezo* in the heart is also essential for courtship competition (Figure 3A). Notably, molecular evidence depicts the importance of *Piezo1* in cardiomyocytes transducing mechanical and chemical signals in mice^46^. Similarly, fruit flies also employ Piezo in the heart to buffer calcium oscillation^47^. Here we have observed that flies with *Piezo* knocked down in the heart show sever defect in mating competition (Figure 3B and Extended Data Figure 6 A, B), suggesting that *Piezo* may transduce mechanical cues from heart contraction to chemical signals, possibly neuropeptides such as Ion transport peptide (ITP).

Next, we used another driver line Tinman-GAL4 which label different cell types of the heart ^48–50^ to knock down Ion transport peptide (ITP), and found they showed defective courtship performance during competition (Figure 3 B).

We then asked whether an elevation of neuropeptide release from the heart can enhance competitiveness in courtship. When ITP was overexpressed in the heart muscle with the TinCΔ4-Gal4(Figure 3 C), male flies won the competition against the control males (Figure 3 D, E).

Analysis of scRNA-seq data from the Fly Cell Atlas^45^ revealed that ITP is widely expressed in heart tissue, with particularly high expression in ostium cells and certain muscle cells (Fig. 3F–G, Extended Data Figure 5 A and 6 C). During courtship, robust ITP expression in these cells is therefore critical for male competitiveness (Fig. 3H).

### Cardiac neuropeptide release enhances courtship competitiveness via TKR99D in P1 neurons

We next investigated whether increased neuropeptide release from the heart can directly enhance courtship behavior. Using an optical pacing assay, we expressed the genetically coded red-shifted activable channel rhodopsin (CsChrimson) in the heart muscle to pace up the heartbeat with light^32^. After stimulated by light pulses, male flies exhibited a stronger competitiveness against the control groups (Fig. 4A–C), indicating that targeted activation of the cardiomyocytes was sufficient to enhance courtship performance in male flies. Notably, the promotion lasted for at least 10 minutes after the light activation, suggesting the cardiac feedback to the brain was mediated through an increase of humoral circulation of certain neuropeptides, likely by ITP.

Neuropeptides are released by secretion of exosomes or dense-core granules (DCGs)^51^. To test whether neuropeptides were indeed released from the heart during courtship, we employed a dense-core granules (DCGs) marker^52^ to monitor the releasing of DCGs from the heart. The DCGs signals decreased dramatically in males’ cardiomyocytes after they were grouped with virgin females for 15 minutes without copulation, implying the vigorous release of ITP containing vesicles from the heart during courtship (Figure 4 D-F).

To investigate how ITP released from the heart promotes sexual behavior, we considered the endocrine system’s capacity for long-range regulation, hypothesizing that circulating ITP might influence neural activity by binding to receptors in the central nervous system. Recently, it’s been identified that ITP could activate TKR99D in flies^53^.To test whether TKR99D mediates ITP’s effects on sexual behavior, we first examined the role of the TKR99D in male courtship behavior, and found that mutation of TKR99D caused severe defect in courtship competition (Figure 4 G).

Since P1 neurons integrate external mating cues and regulate mating state transitions^54^, we hypothesized that cardiac feedback might directly activate this neuronal ensemble. The next question was whether ITP acted on P1 neurons in the brain. It’s been reported that TKR99D is expressed in Fru^M+^ neurons^55^, combined with this we believe that released ITP could directly activates courtship hub-P1 neurons to promote sexual behavior. Mutation of TKR99D impaired male competitiveness (Figure 4G), and knocking down TKR99D in P1 neurons similarly resulted in a severe courtship defect (Figure 4H, K). These data support a feed-forward loop model in which female perception triggers CCAP release and consequent heart rate acceleration, leading to ITP release from cardiomyocytes; the circulating peptides then bind to TKR99D on P1 neurons, sustaining arousal states and enhancing male competitiveness until successful mating (Figure 4I).

### A conserved neuropeptide pathway in mice modulates courtship behavior

To assess whether a similar brain–heart axis exists in mammals, we examined male mice as representative model. Phylogenetic analysis demonstrated that Drosophila TkR99D and its orthologs (including Tac1) in mouse, human, zebrafish, rat, monkey, and cow are evolutionarily conserved (Figure 5A). The TacR1 has been implicated in central nervous system regulation of male mating behavior in mice^56^. In male mice, a 15-minute social interaction with a female resulted in a significant upregulation of Tac1 mRNA in the heart (Figure 5B-C). Using a virus-mediated knockdown approach specifically in cardiomyocytes (Figure 5D,G), we observed that male mice with reduced Tac1 expression displayed decreased overall courtship behaviors, although sniffing latency remained unaffected (Figure 5E). Moreover, knocking down Tac1 in the heart muscles caused a marked delay in ejaculation latency (Figure 5F), indicating the lack of mating arousal because of the absence of cardiac feedback. Pharmacological manipulation of heart rate further corroborated these findings: injection of phenylephrine (PE) decreased courtship behaviors, whereas sodium nitroprusside (SNP) increased heart rate and enhanced courtship activity (Figure 5I–L, Extended Data Figure 7, A-G). These results indicate that, in mice, upregulation of Tac1 in cardiomyocytes upon female stimulation promotes courtship behavior, mirroring the mechanism observed in flies.

**Figure 5.**
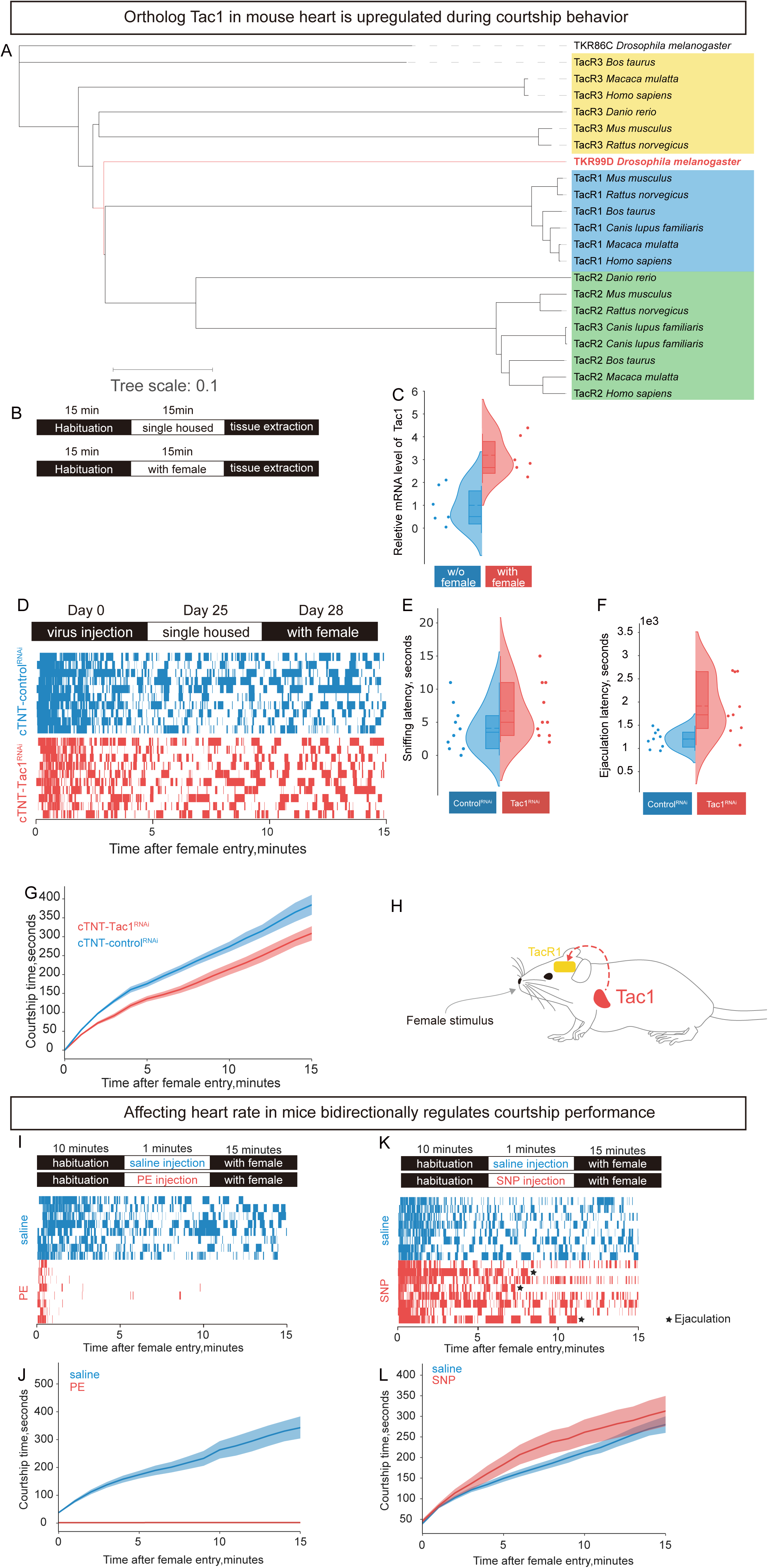
A conserved neuropeptide pathway in mice affects courtship behavior. **A**, Phylogenetic tree showing homology among fly (*Drosophila melanogaster*) TkR99D with mouse (*Mus musculus*), human (*Homo sapiens*), zebrafish (*Danio rerio*), rat (*Rattus norvegicus*), monkey (*Macaca mulatta*) and cow (*Bos taurus*). Here the TKR86C are used as the outer group. **B**, Schematic of the mRNA quantification protocol during male mice social interaction: male mice are habituated in a new cage for 15 minutes before female introduction; after 15 minutes of interaction, males are removed and their hearts are collected for mRNA extraction within 90 minutes. **C**, Male mice exhibit significantly increased *Tac1* mRNA levels in the heart after exposure to female mice. Unpaired t-test, p =0.0011. **D**, Schematic illustrating virus-mediated knockdown of Tac1 specifically in cardiomyocytes and the subsequent behavioral test. Knocking down *Tac1* in heart affected courtship behavior in male mice, all mice without *Tac1* in cardiomyocytes had decreased courtship behavior. **E**, Male mice knocked down *Tac1* in cardiomyocytes exhibit unaffected sniffing latency, indicating the normal mating drive of male mice. Unpaired t-test, p = 0.1570. **F**, Male mice knocked down *Tac1* in cardiomyocytes exhibit longer ejaculation latency, indicating a lower activation at early engagement. Unpaired t-test, p =0.0041. **G**, Cumulatively, male mice with cardiac *Tac1* knockdown display decreased overall courtship time. **H**, Schematic model of the conserved pathway: upon female stimulation, Tac1 expression is upregulated in cardiomyocytes, thereby enhancing male courtship behavior through substance P release. **I-J**, Pharmacological manipulation of heart rate in mice affects courtship behavior. Male mice were firstly introduced to new cage 10 minutes for habituation, then male mice received wither saline or PE before interaction with female mice. Courtship behaviors of male mice were largely decreased after the injection of PE. **K-L**, Same paradigm with SNP injection increased heart rate also enhanced courtship behavior in male mice. Courtship behaviors of male mice were increased after the injection of SNP. Ejaculation events are labeled as asterisk, indicating the hyperactivity of male mice.

### Dynamic regulation of cardiac activity balances competitiveness and long-term fitness

Prolonged exposure to female pheromones without the completion of mating has been shown to have detrimental effects on male flies, including reduced stress resistance and shortened lifespan^57^. We hypothesized that this damage might be tied to the sustained elevation of cardiac activity induced by female cues. In line with this, male flies with an intact cardiac response exhibited diminished resistance to stress and shorter lifespans when exposed to feminized males in the same assay (Figure 6A-B). However, by knocking down CCAP receptors (CCAP-R) specifically on the heart muscle, which prevented the heart rate from accelerating upon female perception, we found that these flies retained normal stress resistance and lifespan (Figure 3B-E). This result indicates that the cardiac response to female cues contributes to long-term physiological change and reduced homeostatic resilience in male flies.

**Figure 6.**
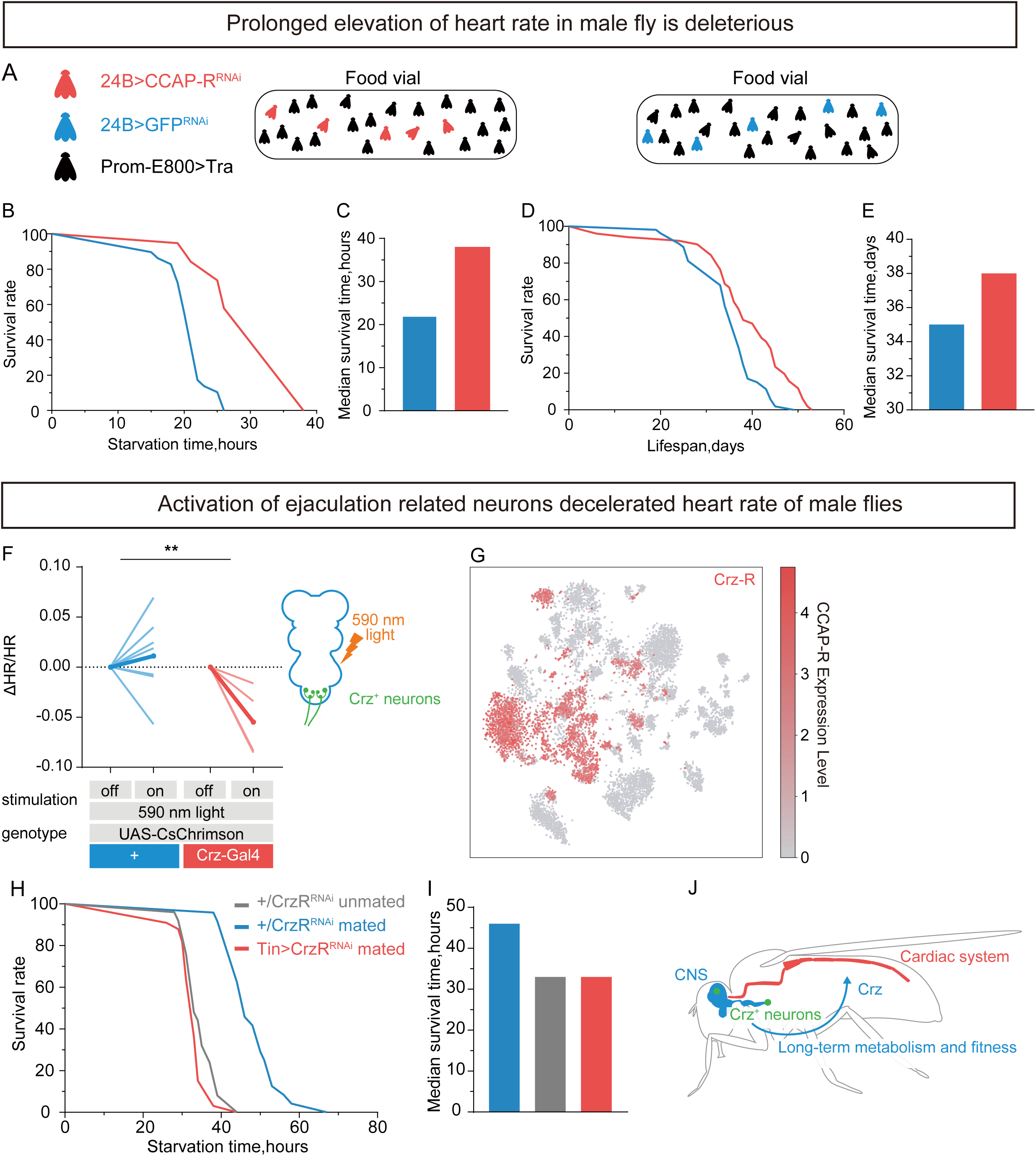
Dynamic regulation of cardiac activity balances transient mating competitiveness and long-term fitness. **A**, Schematic of the female exposure assay: five tester males are grouped with 25 feminized male flies per food vial, based on previous reports that prolonged arousal mating states detrimentally affect male fitness and life span. **B-C**, Cardiac-specific knockdown of the CCAP receptor (CCAP-R) significantly extends the starvation survival time of male flies after female exposure (Log-rank (Mantel-Cox) test, p < 0.0001), indicating that the deleterious effects of female pheromone perception are largely due to prolonged elevated cardiac activity. In this assay, tester flies are placed in empty vials with hydrated filter paper. **D-E**, Similarly, under normal feeding conditions, CCAP-R knockdown in the heart extends male fly lifespan after female exposure (Log-rank test, p < 0.0001), suggesting that sustained cardiac activity during arousal imposes an energetic cost that impairs long-term metabolic fitness. Tester flies are maintained in vials with standard food. **F**, Optogenetic stimulation of ejaculation related Crz^+^ neurons decelerates the heart rate in male flies. Two-way ANOVA, p=0.0024. **G**, The scRNA data revealed the expression pattern of Crz receptors in heart tissue is mainly in pericardial cell and other unannotated cell types, data from fly cell atlas (https://www.flycellatlas.org/). **H-I**, Mating rescues the early death induced by female pheromone exposure (gray and blue). Male flies with CrzR knockdown exhibit reduced starvation resistance (red and blue). Log-rank (Mantel-Cox) test, p<0.0001. **J**, Illustrative schematic depicting the flip of arousal mating state mediated through cardiac system. An unchecked feed-forward loop, in which heart rate remains elevated, is detrimental to fitness and lifespan; however, ejaculation—triggered by Crz^+^ neuron activation—terminates the elevated heart rate and resets the arousal state to a normal resting level, thereby supporting long-term metabolic balance and overall fitness.

Given these harmful effects of sustained cardiac elevation, we next examined whether the natural deceleration of heartbeat after successful mating could mitigate this physiological stress. A success mating is typically terminated with ejaculation^42,58,59^. In male flies, Crz^+^ neurons were reported to coordinate sperm transfer^58^ and ejaculation^59^. We thus hypothesized Crz^+^ neurons decelerated heartbeat after ejaculation. Photoactivation of Crz^+^ neurons can induce ejaculation in male flies^59^. We expressed CsChrimson in Crz^+^ neurons, and found that the heartbeat of the male flies decreased after photoactivation of Crz^+^ neurons (Figure 6 F). Analysis of the RNA sequencing data further confirmed the expression of Crz receptors (CrzR) on the heart, suggesting that Crz released during ejaculation binds to these receptors to calm down the cardiac response (Figure 6 G).

Notably, when Crz receptors were knocked down on the heart, the protective effects of ejaculation on stress resistance were blocked, further underscoring the role of Crz in maintaining post-mating cardiac and physiological stability (Figure 6 H-I).

In summary, these findings reveal that while prolonged cardiac activity due to female cues imposes physiological costs, Crz-mediated deceleration of the heartbeat following ejaculation plays a crucial role in promoting long-term fitness and resilience in male flies, effectively countering the harmful impacts of sustained cardiac elevation (Figure 3 J).

### Cardiac physiology decodes internal mating states flipped by female perception

Based on these molecular and circuitry mechanisms, we developed a novel computational approach to decode internal mating states from cardiac physiology. The cardiac signals were recorded from cardiomyocytes and features extracted via Fourier transform were clustered using dedicated algorithms (Fig. 7A, Extended Data Figure 8). Visualization with an LDA model revealed distinct shifts in the distribution of mating states (Fig. 7B,). The trained decoder achieved a significantly higher area under the ROC curve compared with a decoder trained on shuffled data (Fig. 7C, Extended Data Figure 9). Feature importance analysis via a random forest model and SHAP values identified the most discriminative cardiac parameters (Fig. 7D, E). Finally, distinct stimuli resulted in unique distribution orientations on the LDA plane (Fig. 7F), where internal mating states reflected the unchanged cardiac response (Figure 7 G). These results reflected the cardiac response from distinct stimulation as previously observed (Figure 1 D).

**Figure 7.**
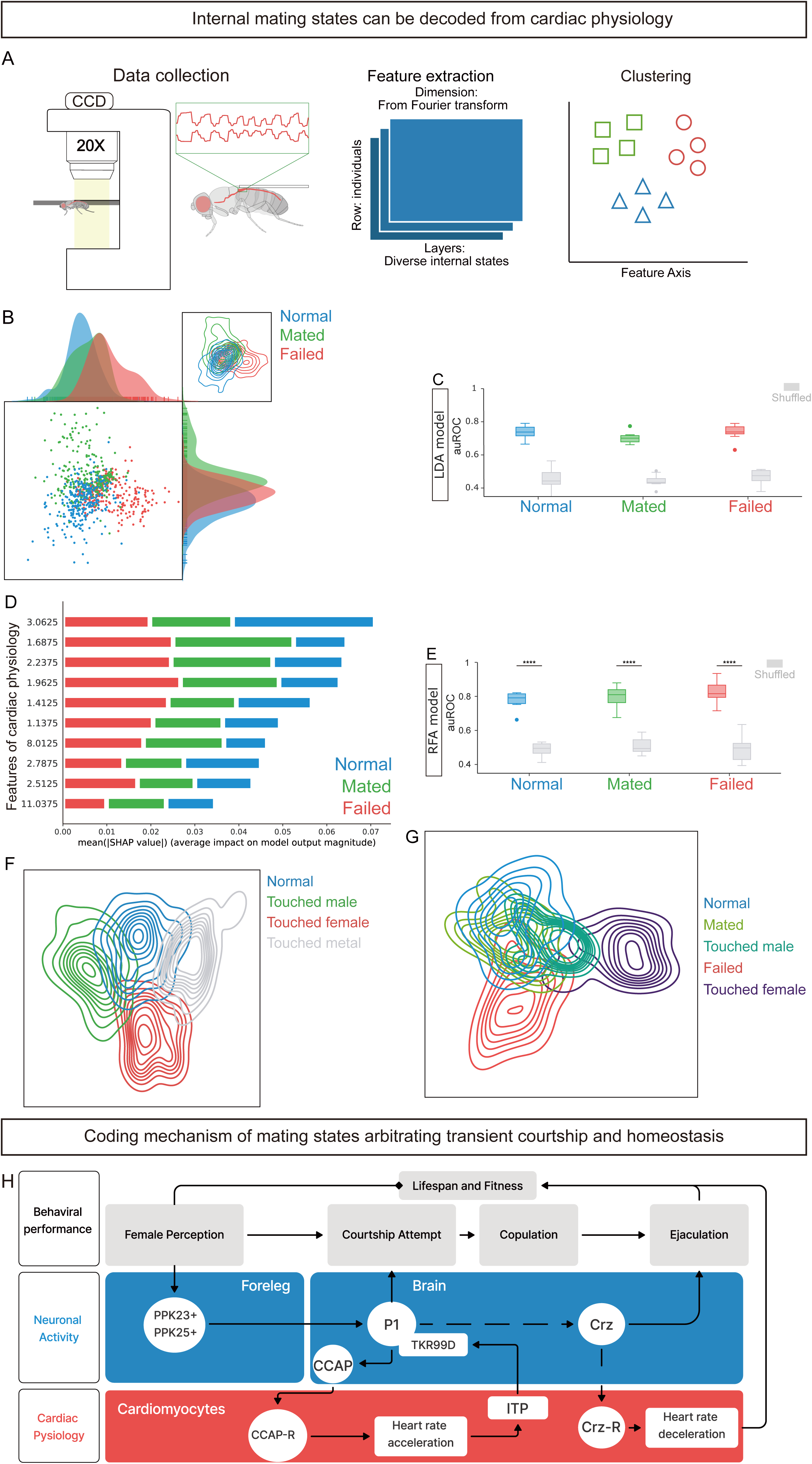
Cardiac physiology decodes internal mating states flipped by female perception. **A**, Schematic illustration of the data processing pipeline: flies are tethered at the posterior abdomen, near the main heart chamber (highlighted in red). Cardiac activities from cardiomyocytes are recorded and features are extracted via Fourier transform for different internal states. These feature matrices are then clustered using dedicated algorithms. **B**, Visualization of high dimensional clustering by LDA model in 2-dimensional plane. Mating states have a shared distribution with separate shifts in such discriminative margin. **C**, The trained decoder in LDA model showed significant higher area under ROC curve(auROC) comparing with shuffled training decoder. Unpaired t-test, p<0.0001. **D**, Most discriminative features are calculated through the random forest model and calculated by SHAP value. **E**, The trained decoder in RFA model showed significant higher area under ROC curve(auROC) comparing with shuffled training decoder. Unpaired t-test, p<0.0001. **F**, Flies subjected to distinct stimuli display separate distribution orientations on the LDA plane. **G**, Mated and normal flies exhibit similar distribution patterns in the LDA plane, whereas failed flies show a distinct separation. Moreover, female stimulation induces a shift in the distribution exclusively in failed flies, while mated flies remain unchanged after perceiving females, consistent with the heart rate changes shown in Figure 1. **H**, Schematic model summarizing the brain–heart communication network in male mating behavior as delineated in this study.

## Discussion

Our study elucidates a previously uncharacterized brain–heart axis that plays a pivotal role in regulating sexual behavior by encoding internal mating states (Figure 7 H and Extend Data Figure 10 A). First, we demonstrated that female perception triggers distinct cardiac activity patterns in male flies and mice. These patterns—reminiscent of the “butterflies in the stomach” phenomenon—serve not only as reflections of internal arousal but also as active drivers of subsequent behavioral execution during courtship. Along with previously studies which have revealed the essential roles of brain-gut, brain-gonad axis in mating behaviors ^15,60^, our study further demonstrates that male animals may synchronize all the organs (here the heart) in order to conduct successful courtship and mating. Furthermore, our results suggest that the brain-heart axis arbitrates the transient challenge and long-term metabolism during mating behaviors. Overall, our study encourages the brain-body network theory in specific behaviors, providing potential opportunities to reveal the evolutionarily conserved connections between innate emotional states and bodily physiology.

### The feed-forward loop of brain-heart axis during courtship behavior

Courtship and copulation are particularly demanding for male animals, requiring them to mobilize their full physiological potential. Our findings suggest that two interlinked mechanisms may underlie this process. First, enhanced cardiovascular function during courtship likely optimizes sensory and motor performance by increasing the delivery of oxygen, nutrients, and hormones to the brain and other critical organs. Second, our data indicate that the cardiac system actively communicates with the central nervous system through neuropeptide signaling. Specifically, during courtship, the heart appears to release neuropeptides that engage the ITP–TKR99D signaling pathway, thereby stimulating P1 neurons in the brain (Figure 4L). This ascending activation creates a feed-forward loop that sustains a heightened internal arousal state and enhances male competitiveness (Extend Data Figure 10 A).

However, several aspects of this model remain to be fully validated. For instance, the precise temporal dynamics of neuropeptide release and signal integration within this feedback loop are not yet clear. It is still uncertain whether the observed positive feedback can be attributed solely to CCAP and ITP signaling, or if additional cardiac-derived signals contribute to the sustained activation of P1 neurons during courtship. Furthermore, dissecting the relative contributions of each component in the loop—such as the exact timing and interplay between CCAP release, ITP secretion, and TKR99D-mediated neural activation—remains an open challenge.

### Cardiac Hyperactivity is terminated after mating to maintain homeostasis

Constant activation of neuronal circuits involved in motivated tasks, without appropriate reward or termination, can jeopardize physiological homeostasis. In fruit flies, continuous activation of female-perception neurons has been associated with early death^57,61^. In mammals, even brief visual or olfactory cues can induce measurable hormonal shifts^62^. While it remains unclear how stress impacts whole-body health, the established link between cardiac function and emotional states^20,22^, suggests that prolonged mental stress may dysregulate cardiovascular health, contributing to systemic decline. Our findings indicate that the brain–heart axis operates not only to boost mating efficacy through transient tachycardia during courtship, but also to safeguard long-term metabolic homeostasis. Once the reproductive demand is satisfied, the system effectively terminates the elevated cardiac state—presumably via regulation through the Crz pathway from nervous system—to return the heart to its resting state. This reset is crucial for preventing the metabolic strain and oxidative stress that could otherwise result from prolonged tachycardia. In doing so, the termination of the feed-forward loop ensures that the benefits of heightened arousal do not come at the expense of overall physiological health.

### Brain-heart axis has conserved functions in sexual behavior across species

On a molecular level, we identified CCAP signaling in flies as a key driver of heart rate acceleration, mediated by CCAP receptors on cardiac muscle^43^ during courtship behavior. Putative analogues of CCAP receptors in mammals, such as oxytocin receptors (OTR) and arginine vasopressin receptors (AVPR1a, AVPR1b)^63^ are known to regulate sexual and social behaviors, as well as heart function. These findings prompt further investigation into whether oxytocin or other neuropeptides are released in response to sexual arousal and modulate heart activity to facilitate mating performance. Additionally, the ITP-TKR99D signaling in flies and Substance P-TacR1 pathway in mice suggest a conserved endocrine feedback loop, bridging cardiac function and central nervous modulation during sexual behavior. Notably, while Substance P plays diverse roles across emotional states and brain regions^9,56^, understanding its function within specific organs and brain circuits remains a key question for future research.

On a circuitry level, cardiac function in mammals is regulated by sympathetic and parasympathetic systems, with recent studies focusing on specific brainstem nuclei and related pathways involved in cardiovascular control ^5,21,64,65^. Meanwhile, cardiac sensory neurons are also being identified ^17,66^. While neural circuits are integral to this system^33^, humoral communication between the brain and heart further diversifies regulatory pathways. Our findings reveal, for the first time, that these pathways contribute to internal mating state encoding through a brain-heart axis, highlighting a previously unexplored role of neuropeptides released from the heart in modulating courtship behavior. The cardiac physiological and behavior results suggest that the mating initiation or initial mating drive may not depend on cardiac activity, as interaction with female mice at the beginning could induce heart beat acceleration and sniffing, and only prolonged elevation of heart rate are required to maintain the persistent courtship behavior. These results together also corroborate the evidence of the heart-derived neuropeptide function in internal mating states in mammals. So, it’s intriguing to validate the function and similar mechanism of brain-heart axis in diverse internal states and other conditions in rodents and primates. However, it remains an open question how the sexual arousal state encoded by the male mammalian brain^56^ is transmitted to the cardiac activity. A comparative interrogation of these mechanisms could provide valuable insights into the molecular and circuit-level interactions of the brain–heart axis in mammalian sexual behaviors, as well as its broader roles in diverse internal states and physiological conditions.

### The central representation of cardiac activity in internal animal states and emotional effects

From a methodological perspective, we have characterized the dynamic regulation of internal state space through three biologically relevant readouts: neuronal dynamics, changes in cardiovascular physiology, and behavioral alterations. By adopting an inter-organ communication perspective, our framework not only delineates how internal states are shaped by context-dependent cardiac activity but also uncovers the encoding mechanisms underlying these interactions. This approach provides a bridge between previous research on neural and behavioral correlates of internal states and a broader physiological framework that incorporates cardiovascular signaling. Our findings open new avenues for investigating how cardiac-derived signals modulate neuronal processes and behavioral outcomes, offering a more integrative model of internal state regulation. By integrating these modalities, future research can explore the precise pathways and interactions involved, potentially leading to novel interventions and therapies targeting the regulation of internal states across various contexts and emotional categories.

## Acknowledgments

We thank the members of the Zhang laboratory for discussions. This work was supported by grants from the Innovation 2030 Major Project of the Ministry of Science and Technology of China (2021ZD0203300). This work was supported by grants 31871059 and 32022029 from the National Natural Science Foundation of China. This work is supported by Chinese Institute for Brain Research, Beijing. W.Z. is an awardee of the Young Thousand Talent Program of China.

## Author contributions

Hongyu Li performed the experiments and analyzed data unless otherwise noted. Shenjie Cheng performed the behavior and survival experiments. Tongxin Diao participated in the optogenetic and immunostaining experiments. Qiruo Zhang performed the surgery of ECG recording assay. Yijun Niu collected the cardiac trace of internal state data. Hongyu Li wrote the manuscript. Wei Zhang supervised the project and edited the manuscript. All authors discussed and commented on the manuscript.

## Data and materials availability

All data needed to evaluate the conclusions in the paper are present in the paper and/or the Supplementary Materials.

## References

1. Anderson, D. J. Circuit modules linking internal states and social behaviour in flies and mice. Nature Reviews Neuroscience 17, 692–704 (2016).

2. Flavell, S. W., Gogolla, N., Lovett-Barron, M. & Zelikowsky, M. The emergence and influence of internal states. Neuron 110, 2545–2570 (2022).

3. Sani, O. G., Abbaspourazad, H., Wong, Y. T., Pesaran, B. & Shanechi, M. M. Modeling behaviorally relevant neural dynamics enabled by preferential subspace identification. Nat. Neurosci. 24, 140–149 (2021).

4. Livneh, Y. & Andermann, M. L. Cellular activity in insular cortex across seconds to hours: Sensations and predictions of bodily states. Neuron 109, 3576–3593 (2021).

5. Chang, R. B., Strochlic, D. E., Williams, E. K., Umans, B. D. & Liberles, S. D. Vagal sensory neuron subtypes that differentially control breathing. Cell 161, 622–633 (2015).

6. Titos, I. et al. A gut-secreted peptide suppresses arousability from sleep. Cell 186, 1382–1397.e21 (2023).

7. Wang, P., Jia, Y., Liu, T., Jan, Y. N. & Zhang, W. Visceral Mechano-sensing Neurons Control Drosophila Feeding by Using Piezo as a Sensor. Neuron 108, 640–650.e4 (2020).

8. Yang, D. et al. Nociceptor neurons direct goblet cells via a CGRP-RAMP1 axis to drive mucus production and gut barrier protection. Cell 185, 4190–4205.e25 (2022).

9. Zhang, W. et al. Gut-innervating nociceptors regulate the intestinal microbiota to promote tissue protection. Cell 185, 4170–4189.e20 (2022).

10. Bin, N. R. et al. An airway-to-brain sensory pathway mediates influenza-induced sickness. Nature 615, (2023).

11. Tao, J. et al. Highly selective brain-to-gut communication via genetically defined vagus neurons. Neuron 109, 2106–2115.e4 (2021).

12. Lin, H. H. et al. A nutrient-specific gut hormone arbitrates between courtship and feeding. Nature 602, 632–638 (2022).

13. Ahmed, S. M. H. et al. Fitness trade-offs incurred by ovary-to-gut steroid signalling in Drosophila. Nature 584, 415–419 (2020).

14. Hadjieconomou, D. et al. Enteric neurons increase maternal food intake during reproduction. Nature 587, 455–459 (2020).

15. Hudry, B. et al. Sex Differences in Intestinal Carbohydrate Metabolism Promote Food Intake and Sperm Maturation. Cell 178, 901–918 (2019).

16. Zeng, W. Z. et al. PIEZOs mediate neuronal sensing of blood pressure and the baroreceptor reflex. Science *(80-.).* 362, 464–467 (2018).

17. Salameh, L. J., Bitzenhofer, S. H., Hanganu-Opatz, I. L., Dutschmann, M. & Egger, V. Blood pressure pulsations modulate central neuronal activity via mechanosensitive ion channels. Science (80-.). 383, (2024).

18. Klein, A. S., Dolensek, N., Weiand, C. & Gogolla, N. Fear balance is maintained by bodily feedback to the insular cortex in mice. Science *(80-.).* 374, 1010–1015 (2021).

19. Barrios, N., Farias, M. & Moita, M. A. %J A. at S. 3733187. Threat induces cardiac and metabolic changes that negatively impact survival in flie. Curr. Biol. (2020). 10.1016/j.cub.2021.10.013

20. Nummenmaa, L., Glerean, E., Hari, R. & Hietanen, J. K. Bodily maps of emotions. Proc. Natl. Acad. Sci. U. S. A. 111, 646–651 (2014).

21. Critchley, H. D. & Harrison, N. A. Visceral Influences on Brain and Behavior. Neuron 77, 624–638 (2013).

22. Rainville, P., Bechara, A., Naqvi, N. & Damasio, A. R. Basic emotions are associated with distinct patterns of cardiorespiratory activity. Int. J. Psychophysiol. 61, 5–18 (2006).

23. White, G. L., Fishbein, S. & Rutsein, J. Passionate love and the misattribution of arousal. J. Pers. Soc. Psychol. 41, 56–62 (1981).

24. Zhao, B. et al. Heart-brain connections: Phenotypic and genetic insights from magnetic resonance images. Science *(80-.).* 380, abn6598 (2023).

25. Resch, J. et al. Neural basis for fasting activation of the hypothalamic–pituitary–adrenal axis. Nature 620, 154–162 (2023).

26. Kostis, J. B. et al. Sexual dysfunction and cardiac risk (the Second Princeton Consensus Conference). Am. J. Cardiol. 96, 313–321 (2005).

27. Litviňuková, M. et al. Cells of the adult human heart. Nature 588, 466–472 (2020).

28. Huynh, P. et al. Myocardial infarction augments sleep to limit cardiac inflammation and damage. Nature 635, (2024).

29. Ziegler, K. A. et al. Immune-mediated denervation of the pineal gland underlies sleep disturbance in cardiac disease. Science (80-.). 381, 285–290 (2023).

30. Wagner, J. U. G. G. et al. Aging impairs the neurovascular interface in the heart. Science (80-.). 381, 897–906 (2023).

31. Skelly, D. A. et al. Single-Cell Transcriptional Profiling Reveals Cellular Diversity and Intercommunication in the Mouse Heart. Cell Rep. 22, 600–610 (2018).

32. Hsueh, B. et al. Cardiogenic control of affective behavioural state. Nature 615, 292–299 (2023).

33. Zhao, Q. et al. A multidimensional coding architecture of the vagal interoceptive system. Nature 603, 878–884 (2022).

34. Prochazkova, E., Sjak-Shie, E., Behrens, F., Lindh, D. & Kret, M. E. Physiological synchrony is associated with attraction in a blind date setting. *Nat*. Hum. Behav. 6, 269–278 (2022).

35. Barske, J., Schlinger, B. A., Wikelski, M. & Fusani, L. Female choice for male motor skills. Proc. R. Soc. B Biol. Sci. 278, 3523–3528 (2011).

36. Orihuela, A., Omaña, J. C. & Ungerfeld, R. Heart rate patterns during courtship and mating in rams and in estrous and nonestrous ewes (Ovis aries). J. Anim. Sci. 94, 556–562 (2016).

37. Terada, M. et al. Changes in the heart rate and plasma epinephrine and norepinephrine concentrations of the stallion during copulation. Reprod. Med. Biol. 4, 143–147 (2005).

38. Gerber, P. & Schnell, C. R. Behavioral and cardiophysiological responses of common marmosets ( Callithrix jacchus) to confrontations with opposite-sexed strangers. Primates 45, 187–196 (2004).

39. Thistle, R., Cameron, P., Ghorayshi, A., Dennison, L. & Scott, K. Contact chemoreceptors mediate male-male repulsion and male-female attraction during drosophila courtship. Cell 149, 1140–1151 (2012).

40. Kallman, B. R., Kim, H. & Scott, K. Excitation and inhibition onto central courtship neurons biases drosophila mate choice. Elife 4, e11188 (2015).

41. Ribeiro, I. M. A. et al. Visual Projection Neurons Mediating Directed Courtship in Drosophila. Cell 174, 607–621 (2018).

42. Yamamoto, D. & Koganezawa, M. Genes and circuits of courtship behaviour in Drosophila males. Nat. Rev. Neurosci. 14, 681–692 (2013).

43. Dulcis, D., Levine, R. B. & Ewer, J. Role of the neuropeptide CCAP in Drosophila cardiac function. J. Neurobiol. 64, 259–274 (2005).

44. Mandal, L., Banerjee, U. & Hartenstein, V. Evidence for a fruit fly hemangioblast and similarities between lymph-gland hematopoiesis in fruit fly and mammal aorta-gonadal-mesonephros mesoderm. Nat. Genet. 36, 1019–1023 (2004).

45. Li, H. et al. Fly Cell Atlas: A single-nucleus transcriptomic atlas of the adult fruit fly. Science (80-.). 375, eabk2432 (2022).

46. Jiang, F. et al. The mechanosensitive Piezo1 channel mediates heart mechano-chemo transduction. Nat. Commun. 12, 896 (2021).

47. Zechini, L. et al. Piezo buffers mechanical stress via modulation of intracellular Ca2+ handling in the Drosophila heart. Front. Physiol. 13, 1–34 (2022).

48. Wessells, R. J., Fitzgerald, E., Cypser, J. R., Tatar, M. & Bodmer, R. Insulin regulation of heart function in aging fruit flies. Nat. Genet. 36, 1275–1281 (2004).

49. Han, Z., Yi, P., Li, X. & Olson, E. N. Hand, an evolutionarily conserved bHLH transcription factor required for Drosophila cardiogenesis and hematopoiesis. Development 133, 1175–1182 (2006).

50. Lo, P. C. H. & Frasch, M. A role for the COUP-TF-related gene seven-up in the diversification of cardioblast identities in the dorsal vessel of Drosophila. Mech. Dev. 104, 49–60 (2001).

51. Pols, M. S. & Klumperman, J. Trafficking and function of the tetraspanin CD63. Exp. Cell Res. 315, 1584–1592 (2009).

52. Redhai, S. et al. Regulation of Dense-Core Granule Replenishment by Autocrine BMP Signalling in Drosophila Secondary Cells. PLoS Genet. 12, 1–32 (2016).

53. Xu, W., Li, G., Chen, Y., Ye, X. & Song, W. A novel antidiuretic hormone governs tumour-induced renal dysfunction. Nature 624, 425–432 (2023).

54. Jung, Y. et al. Neurons that Function within an Integrator to Promote a Persistent Behavioral State in Drosophila. Neuron 105, 322–333.e5 (2020).

55. Wohl, M. P., Liu, J. & Asahina, K. Drosophila Tachykininergic Neurons Modulate the Activity of Two Groups of Receptor-Expressing Neurons to Regulate Aggressive Tone. J. Neurosci. 43, 3393–3420 (2023).

56. Bayless, D. W. et al. A neural circuit for male sexual behavior and reward. Cell 186, 3862–3881.e28 (2023).

57. Maures, T. J. et al. Drosophila Life Span and Physiology Are Modulated by Sexual Perception and Reward. Science (80-.). 343, 544–548 (2014).

58. Tayler, T. D., Pacheco, D. A., Hergarden, A. C., Murthy, M. & Anderson, D. J. A neuropeptide circuit that coordinates sperm transfer and copulation duration in Drosophila. Proc. Natl. Acad. Sci. U. S. A. 109, 20697–20702 (2012).

59. Zer-Krispil, S. et al. Ejaculation Induced by the Activation of Crz Neurons Is Rewarding to Drosophila Males. Curr. Biol. 28, 1445–1452 (2018).

60. Redhai, S. et al. An intestinal zinc sensor regulates food intake and developmental growth. Nature 580, 263–268 (2020).

61. Shohat-Ophir, G., Kaun, K. R., Azanchi, R. & Heberlein, U. Sexual deprivation increases ethanol intake in Drosophila. Science (80-.). 335, 1351–1355 (2012).

62. Arrieta, J. M. et al. Oxytocin-gaze positive loop and the coevolution ofhuman-dog bonds. Science (80-.). 348, 331–333 (2015).

63. Kriventseva, E. V. et al. OrthoDB v10: Sampling the diversity of animal, plant, fungal, protist, bacterial and viral genomes for evolutionary and functional annotations of orthologs. Nucleic Acids Res. 47, D807–D811 (2019).

64. Yao, Y. et al. Cardiovascular baroreflex circuit moonlights in sleep control. Neuron 110, 3986–3999 (2022).

65. Prescott, S. L. & Liberles, S. D. Internal senses of the vagus nerve. Neuron 110, 579–599 (2022).

66. Veerakumar, A., Yung, A. R., Liu, Y. & Krasnow, M. A. Molecularly defined circuits for cardiovascular and cardiopulmonary control. Nature 606, 739–746 (2022).

