## Supplemental Figure for "A heart releasing neuropeptide that synchronizes brain-heart regulation during courtship behavior"

**Supplemental Figures**

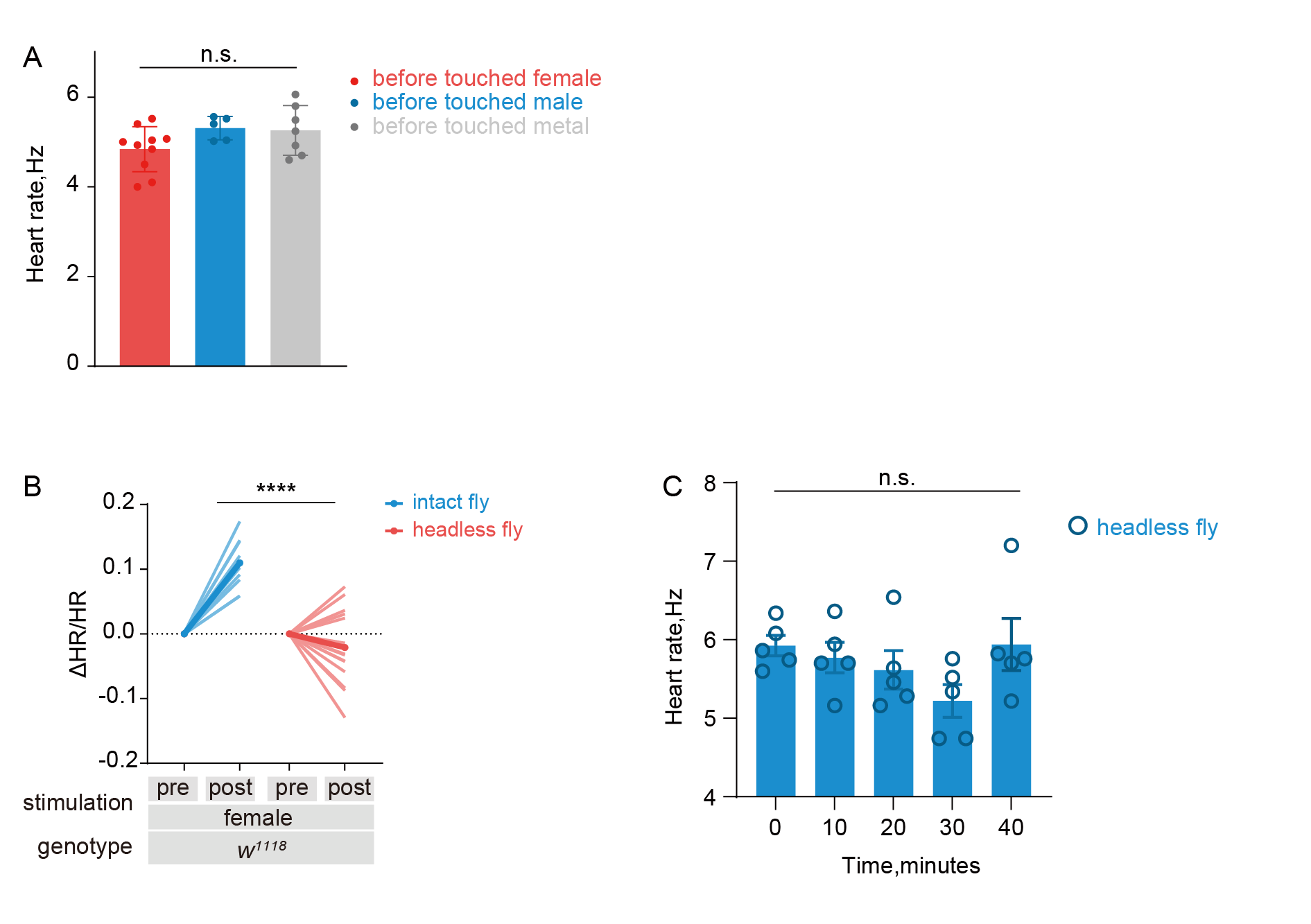

**Extend Data Figure1. The cardiac response of male flies is specialized for courtship behavior through central nervous system**

A, heart rate of male flies is at same level before any external stimulation. Two-way analysis of variance (ANOVA) followed by Tukey’s test, p=0.1923.

B, The cardiac activity was not elevated in decapitated male flies to female stimulation. Two-way analysis of variance (ANOVA) followed by Tukey’s test, p<0.0001.

C, A decapitated male fly exhibits stable heart rate up to 40 minutes.

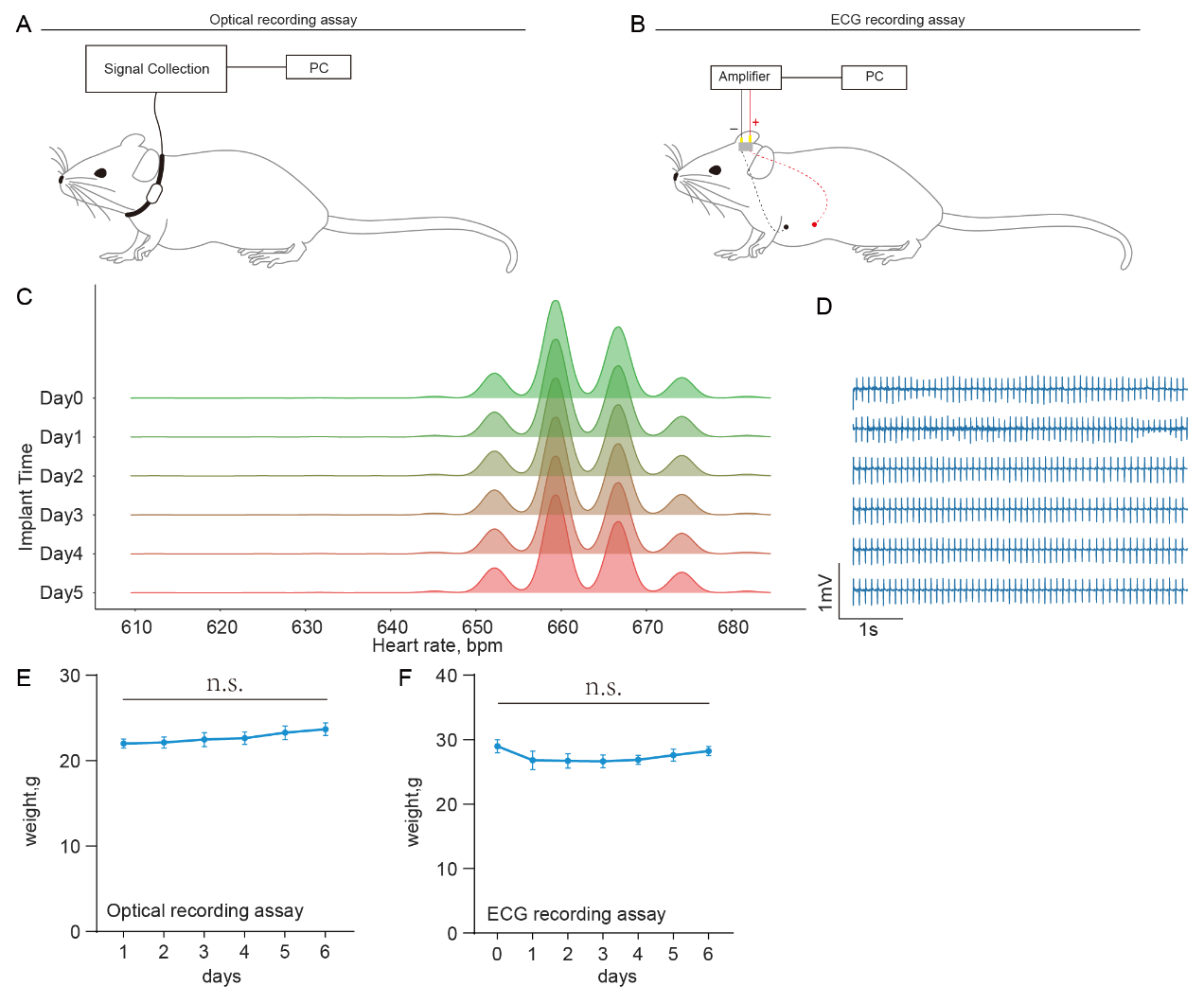

**Extend Data Figure2. The recording assay of heart rate in mice**

A, Illustrative figure of optical recording assay.

B, Illustrative figure of ECG recording assay.

C, Representative heart rate distribution from ECG recording after electrode implant from day 0 to day 5.

D, Representative ECG signals from C.

E, Weight measured each day after habituation under optical recording assay.

F, Weight measured each day after electrode implant in the ECG recording assay.

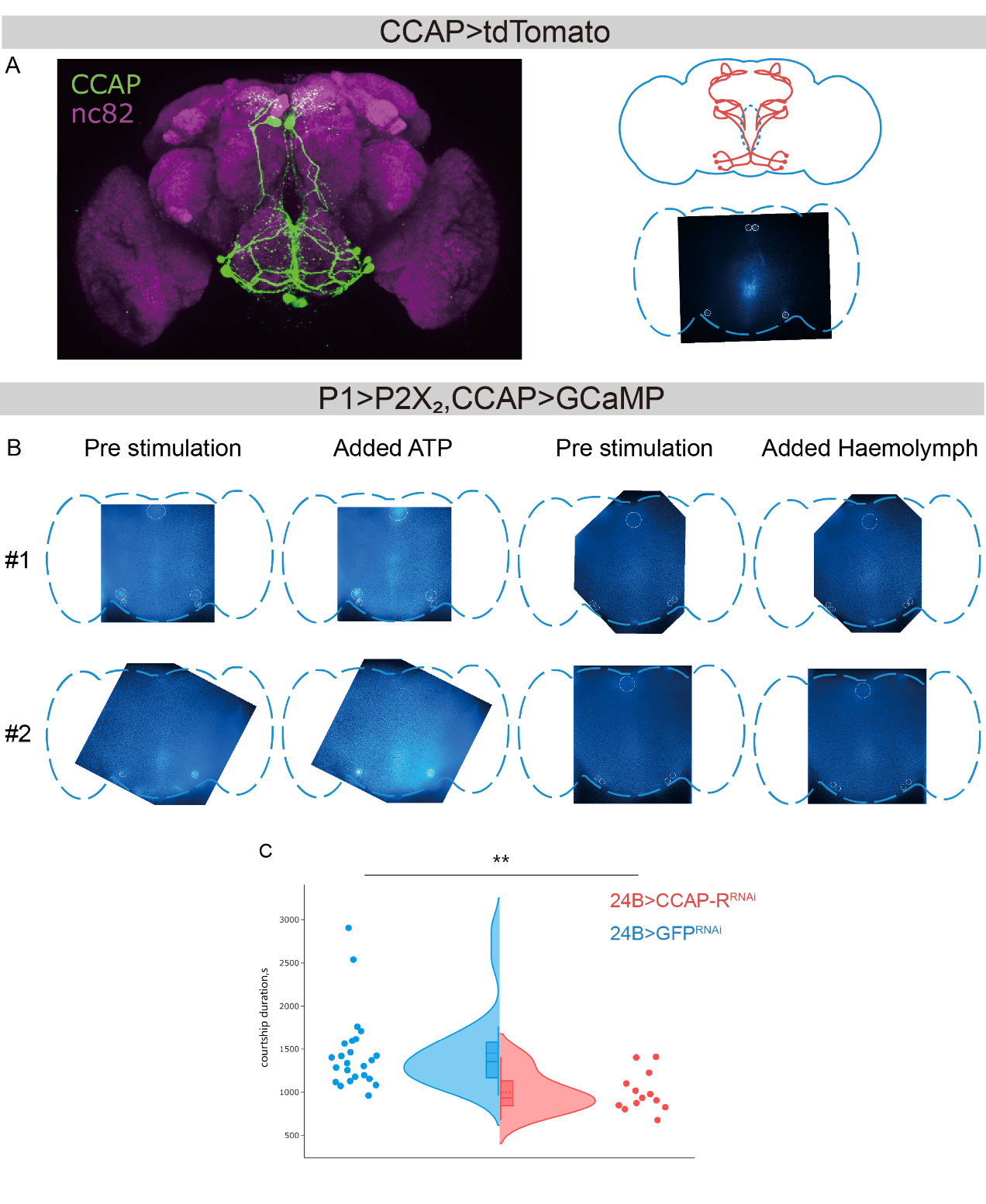

**Extend Data Figure3. P1 triggers CCAP release through activating CCAP^+^ neurons in brain**

A, Expression pattern of CCAP-Gal4 in the brain. Right panel indicated the ROI of calcium imaging for each CCAP neurons in white dashed circle.

B, Representative calcium imaging results. The fluorescence of CCAP neurons is elevated after adding ATP to activate P1 neurons. ROIs are indicated as white dashed circle.

D, Knocking down CCAP-R in the heart muscle affects courtship duration but not courtship latency.

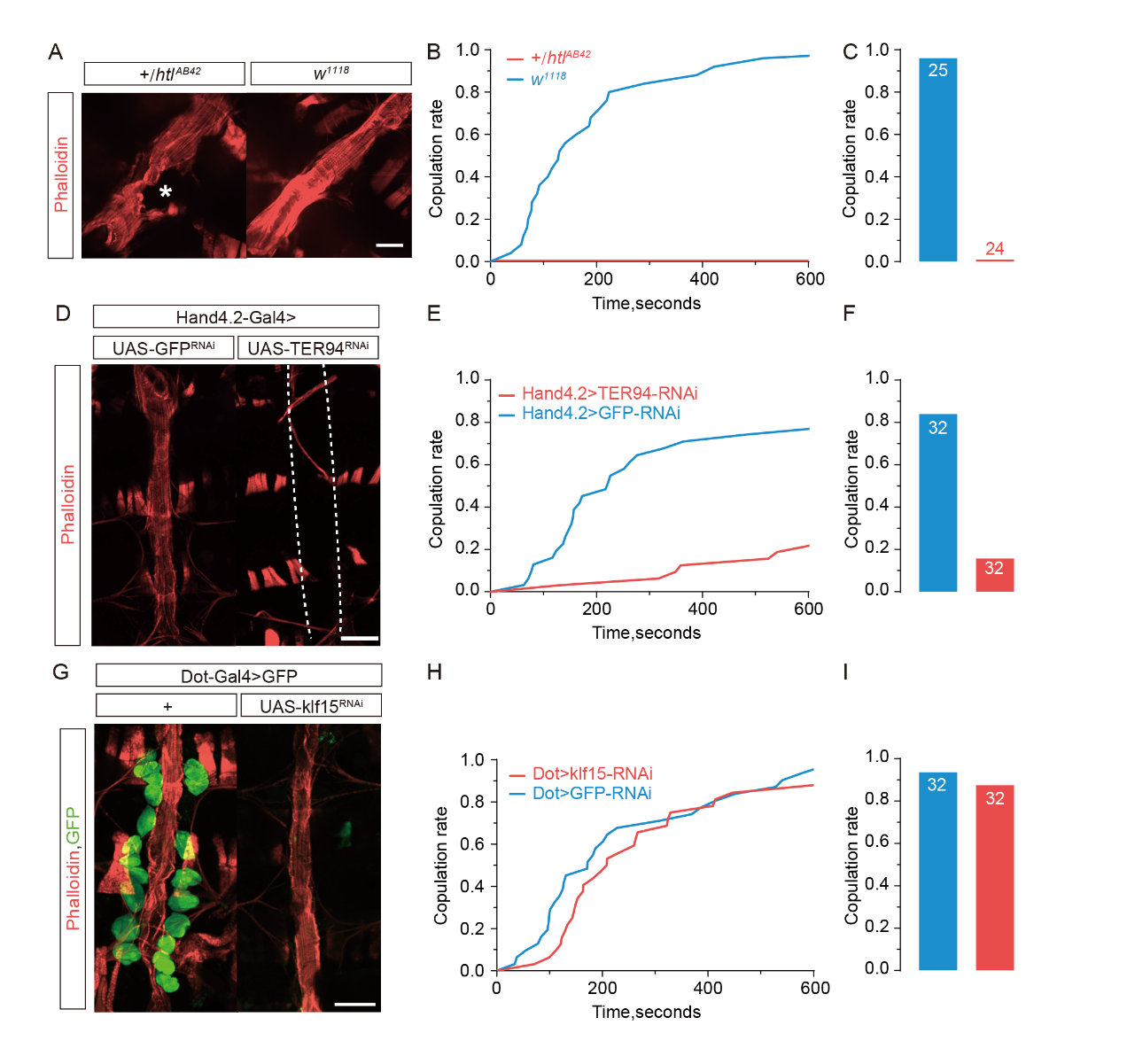

**Extend Data Figure4. The cardiac system of male flies is essential for courtship behavior**

A, Representative images of the heart in *htl* mutant flies and *w^1118^* flies. *Htl* mutation causes severe defect of the integrity of the heart tube (highlighted with asterisk). Scale bar represents 50 μm.

B-C, *Htl* flies exhibit impaired courtship performance toward female flies. Replicates are indicated with numbers on the bars, accumulative success rates were compared by log-rank (Mantel-Cox) test p<0.0001.

D, The genetic approach specifically abolishes the heart tube formation during development. White dashed lines outline the lost heart tube. Scale bar represents 100 μm.

E-F, Male flies lacking the heart tube failed to conduct proper courtship behavior. Replicates are indicated as numbers on the bars, accumulative success rates are compared by log-rank (Mantel-Cox) test p<0.0001.

G, The genetic approach specifically eliminates the pericardial cells surrounding the heart tube. Scale bar represents 100 μm.

H-I, Male flies without the pericardial cells exhibit normal courtship behavior. Replicates are indicated as numbers on the bars, accumulative success rates were compared by log-rank (Mantel-Cox) test, p=0.2026.

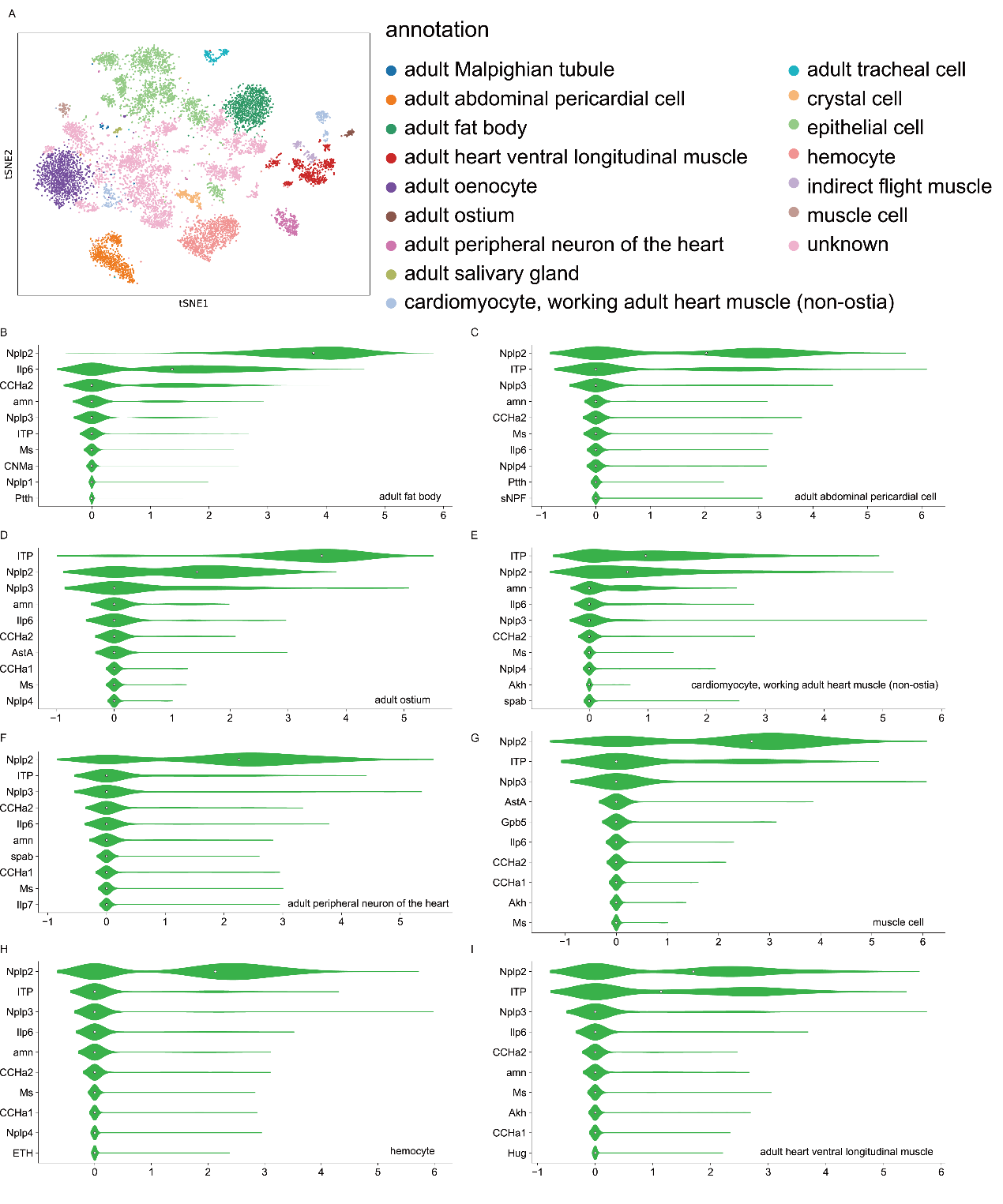

**Extend Data Figure5. Identification of ligand and receptor genes from single cell RNA sequencing data in cardiac system**

A, The RNA sequencing data of the heart tissue clustered by annotation and represented with the tSNE method.

B-I, Expression level of top 10 ligand genes in the heart and associated tissues (fat body, abdominal pericardial cell, ostium, cardiomyocytes, heart muscle, hemocyte, peripheral neurons, ventral longitudinal muscle, muscle cells).

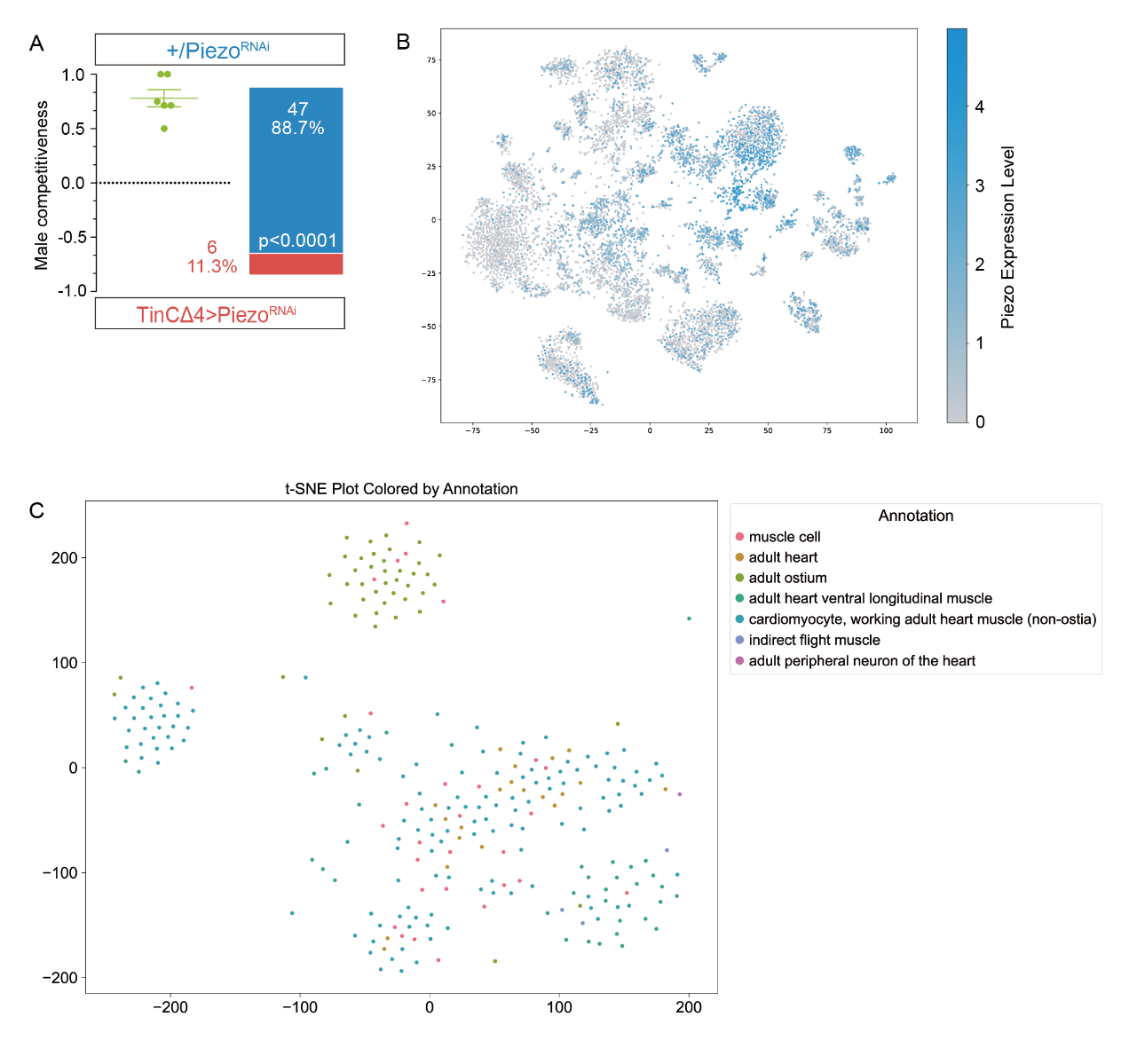

**Extend Data Figure 6. Cardiac signals affect courtship competition**

A, Knocking down Piezo in the cardiomyocyte impairs male competitiveness. Binomial test, p value is noted in figure.

B, Piezo is widely expressed in unannotated cell types, data is from 10X heart tissue(<https://www.flycellatlas.org/>)

C, detailed annotated cardial cells reveals more specific gene expression data is from 10X cardial cell (<https://www.flycellatlas.org/>).

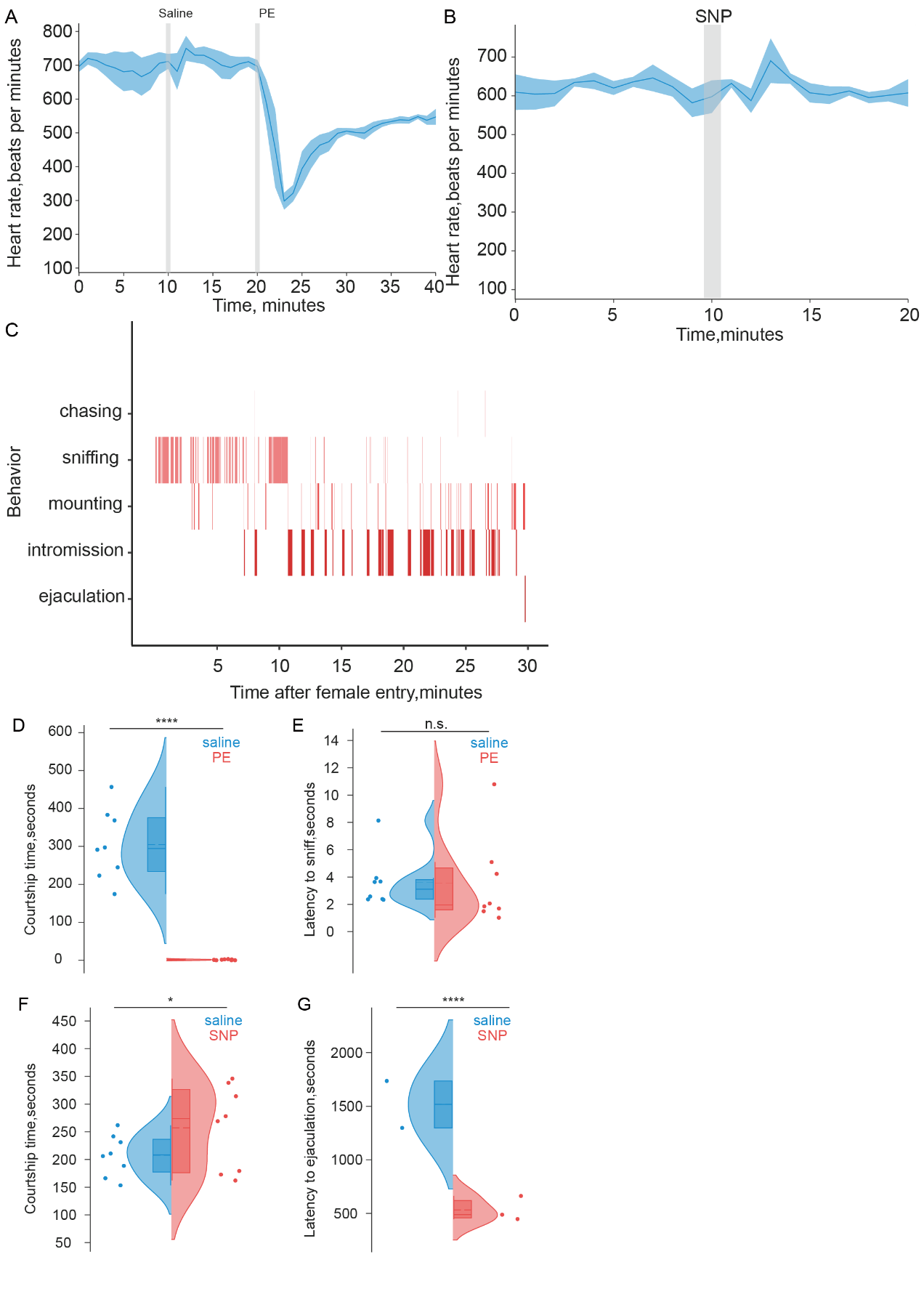

**Extend Data Figure 7. Manipulation of heart rate in male mice largely affects courtship performance**

A, Cardiac response of saline and PE injection, after injection of PE, heart rate of mice was decreased transiently and maintained at low level for more than 20 minutes, heart rate was recorded by ECG recording assay.

B, Cardiac response of saline and SNP injection, after injection of SNP, heart rate of mice was increased transiently and maintained for very short period, heart rate was recorded by ECG recording assay.

C, Representative behavioral results of one individual mouse during courtship.

D, Statistical comparison of accumulative courtship time of male mice receiving either saline or PE. Paired t-test, p <0.0001.

E, Male mice received PE or saline exhibit similar latency to sniffing behavior, indicating the same mating motivation at early engagement. Unpaired t-test, p = 0.9423.

F, Statistical comparison of accumulative courtship time of male mice receiving either saline or SNP. Paired t-test, p =0.04.

G, Male mice received SNP exhibit lower latency to ejaculation behavior, indicating the hyperactivity of male mice at early engagement. Unpaired t-test, p <0.0001.

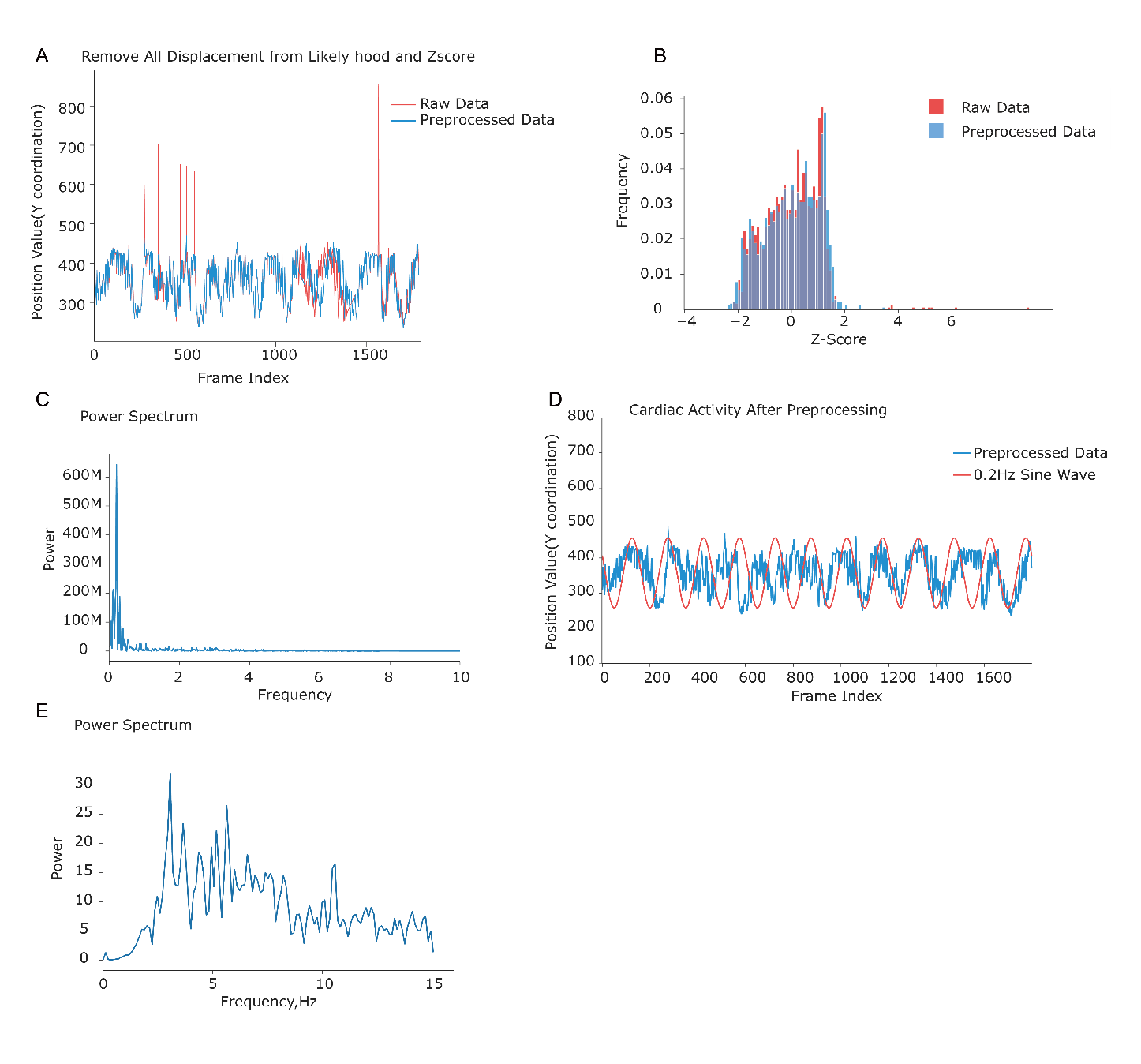

**Extend Data Figure 8. Feature extraction of cardiac physiology of fruit flies.**

A, Preprocessed raw signals are then validated by removing data below a given precision 0.9 (provided by DeepLabCut https://www.mackenziemathislab.org/deeplabcut).

B, Raw data are then performed with Z-score normalization, eliminating data exceeding a given standard deviation(sigma<3).

C, The PSD analysis provides valuable insights into the frequency characteristics of the cardiac signal, including the main frequency and overall frequency distribution.

D, A direct moving artifacts at 0.2 Hz is observed to disturb the real cardiac activity traces.

E, Final cardiac signals with a high-pass filter can be applied for further feature extraction.

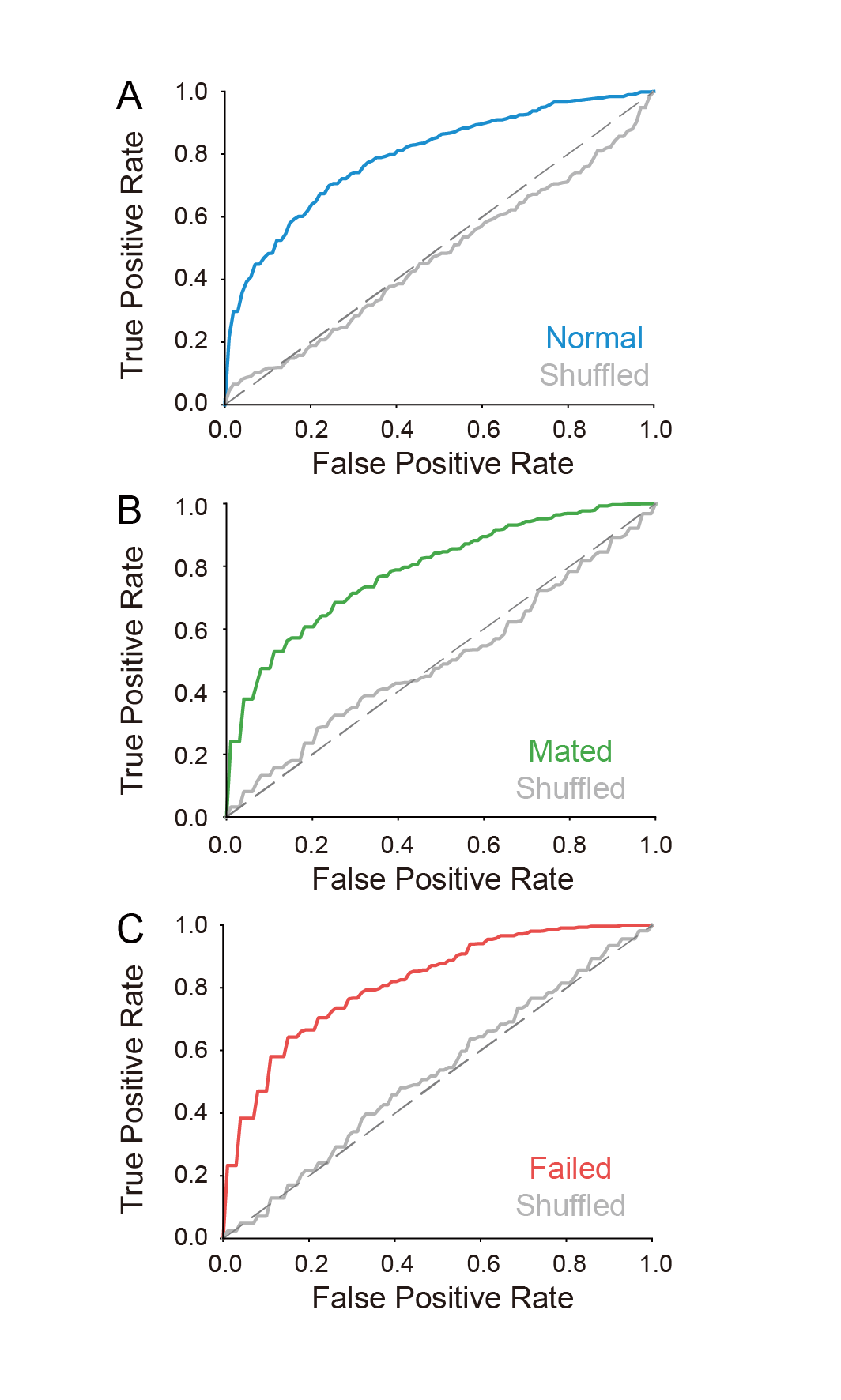

**Extend Data Figure9. ROC curve for mating state decoder by LDA model**

A, Normal mating state can be decoded with trained decoder with a high discriminative rate.

B, Mated state can be decoded with trained decoder with a high discriminative rate.

C, Failed mating state can be decoded with trained decoder with a high discriminative rate.

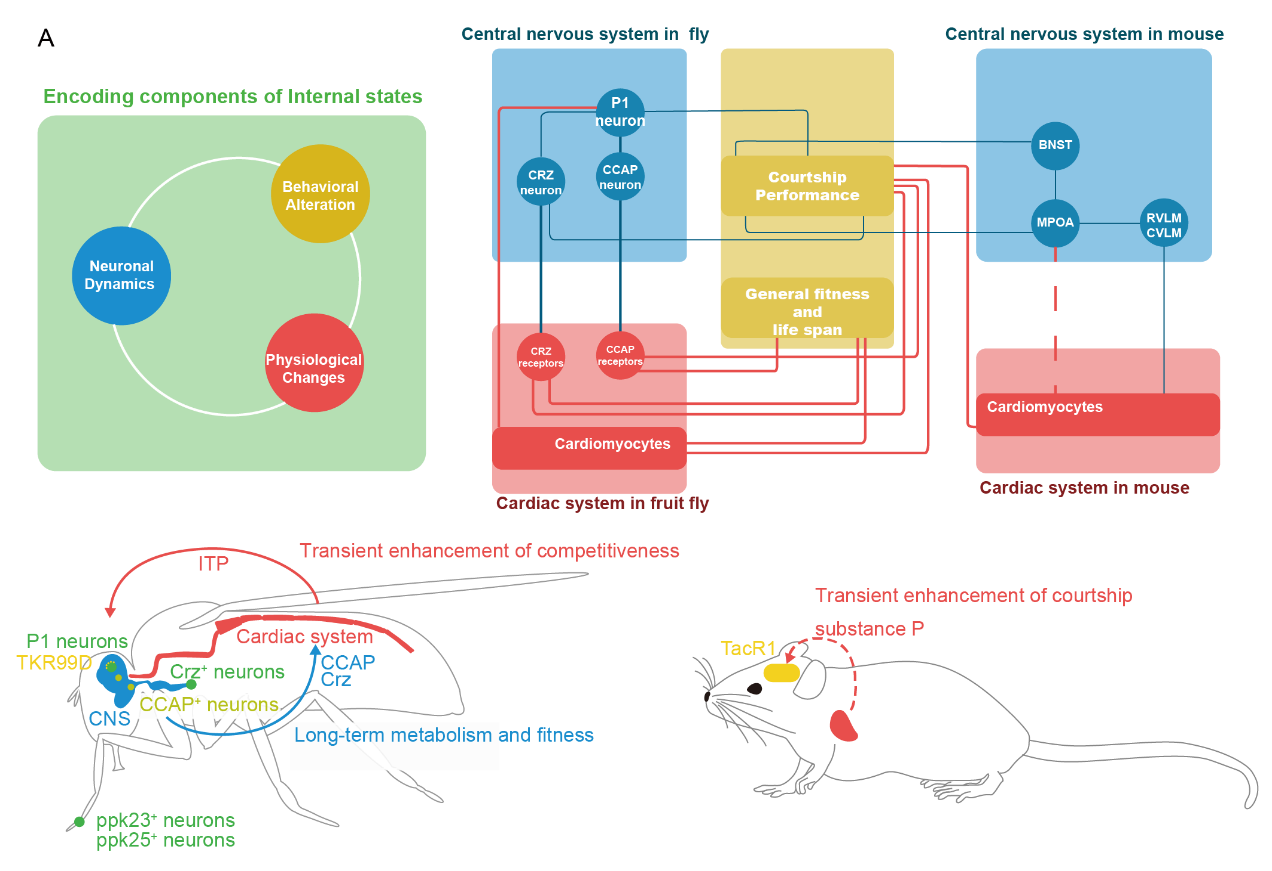

**Extend Data Figure 10. The brain-heart axis in fruit fly and mice shared similar molecular pathway**

A, Illustrative figure of the encoding components of internal states. The brain-heart axis synchronized mating internal states through similar endocrine pathway with conserved molecules.
