## Supplementary Information for "A heart releasing neuropeptide that synchronizes brain-heart regulation during courtship behavior"

**Materials and method**

**Fly husbandry**

All strains (supplementary tables) were maintained on standard cornmeal and sugar media in a 25℃ incubator with 12-hour light/dark cycles unless otherwise noted. Male flies were separated and singly housed within 2 hours after eclosion.

All courtship behaviors were carried out with 5-12 day old sexually mature flies and were video-recorded using cameras. Flies were marked one day before the day of the behavioral assays. Flies were aspirated from one chamber to another unless otherwise noted. For every experiment described below when assays were performed over multiple days, control and experimental animals were tested on the same day.

**Mice**

C57BL/6J mice were obtained from Laboratory Animal Resources Center, Tsinghua University and were housed in groups under standard conditions according to the Tsinghua University animal facility. Mice were maintained on a 12-h light/dark cycle and tested during the light phase of the cycle. All animal work complied with ethical regulations for animal testing and research, and was done in accordance with Institutional Animal Care and Use Committee approval by Tsinghua University and followed all Association for Assessment and Accreditation of Laboratory Animal Care guidelines.

**Flies**

**Video recording of the cardiac activity**

All flies were raised on standard food for 3-5 days after eclosion. A male fly was briefly anesthetized on ice and glued on its back to a 24×24 mm coverslip by Norland Optical adhesive #61 cured using a 365 nm 10-watt UV LED for 20 seconds. Tethered flies were allowed to recover for 10 minutes under the imaging microscope before recording. The coverslip was mounted to the microscope using an assembly containing a small Micro operation platform, which allowed the recording of each heart chamber to optimized. All recording were done under 25℃ and dark condition unless otherwise noted. All tester female flies were newly anesthetized for each day and only their abdominal segments were approached to touch the male fly foreleg.

**Heartbeat quantification**

All recorded videos were converted into MP4 format of 60-second length, and then imported into the ImageJ software. Videos for each individual flies were digitalized using the time series analyzer plugin by the same ROI. The measured intensity data were saved in CSV format. All saved CSV files were imported into MATLAB to denoise the heartbeat signal. Heart rate calculation was conducted in a custom-made MATLAB script (supplementary information). All changes of heart rate were calculated as following:

$$\Delta HB(t)=\frac{HB\left( t \right)-HB(t_{0})}{HB(t_{0})}$$

, where $HB\left( t \right)$ is the heartbeat rate at the recorded time point, $HB(t_{0})$ is the normal heartbeat rate after tethering and recovery before any external stimulation.

**Measuring internal heart states**

Flies during different internal states were raised under same 12-hour light/dark cycles. Male flies were separated and singly housed within 2 hours after eclosion. For aging group, all male flies were single housed after eclosion for 6, 14 or 28 days. Food vials were replaced every 3 days. For different nutritional states, all flies used were 6-8 days old, flies in thirst group were transferred into empty vials 12-15 hours before recording; flies in the starved group were transferred to new vials containing 1% agar for water supplies 12-15 hours before recording. For mating group, all male flies were introduced to a similar chamber as in the courtship competition assay with a *w^1118^* virgin female fly aged 4 to 6 days.

- Raw data acquisition: As we introduced in main results, our data is collected by same recording method (Video recording of the cardiac activity). All recorded videos were converted into MP4 format of 60-second, tracing data from DeepLabCut^TM^ (DLC)^1^ with minimum labeling frame, typically 50 frames for each video (Supplemental video 1).
- Preprocessing: Initially, we preprocessed the raw signals by removing data below a given precision 0.9 (provided by DeepLabCut^TM^ (DLC)^1^), performing Z-score normalization, eliminating data exceeding a given standard deviation(sigma<3), and handling NaN values. This step aids in noise and outlier removal, enhancing the accuracy of subsequent analyses. A representative figure can illustrate the signal before and after preprocessing (Extended Data Figure 8 B).
- Power Spectral Density (PSD): The PSD analysis provides valuable insights into the frequency characteristics of the cardiac signal, including the main frequency and overall frequency distribution. In this step, we applied a high-pass filter (cutoff frequency > 1 Hz) to eliminate motion artifacts from tethered flies, ensuring that the resulting frequency analysis accurately reflects true cardiac activity (Extended Data Figure 8 C-E).

The prepared data was standardized and saved as a data matrix, labeled with the corresponding animal condition for subsequent use in decoder construction. The fast Fourier transform was conducted through scipy^2^.

- Linear discriminative algorithm (LDA) model training

Briefly, all collected data with corresponding state label is applied for construction of decoder. Training was conducted through sklearn with balanced data category (for each group, “normal” labels are more than other internal states). The F1 score and accuracy of training and test were calculated as average for 100 random epochs (Figure). The area under the Receiver Operating Characteristic (auROC) curve was calculated using the trained decoder, both with correct labels and shuffled labels, employing 10-fold cross-validation.

- Gaussian Mixture Model (GMM)

A Gaussian Mixture Model (GMM) was applied as an additional indicator to validate our Linear Discriminant Analysis (LDA) model. We calculated the nearest centroid for each GMM cluster and mapped these clusters to the corresponding LDA labels. This mapping allowed us to assess the true distribution of cardiac activity associated with specific internal states, thereby providing insights into the relationship between the two classification methods.

- Random Forest model

We applied the Random Forest algorithm to further investigate the classification of cardiac activity data across distinct internal heart states. The contribution of each feature was evaluated using SHAP values, providing a deeper understanding of the factors driving classification.

**Fly Courtship Competition assay**

Courtship competition assay was carried out in a 12mm diameter chamber lined with a piece of wet filter paper at 25℃. All *w^1118^* female flies were collected within 2 hours after eclosion and housed in group. Male flies were collected and singly housed right after eclosion. Female flies aged for 4-6 days were briefly anesthetized on ice and introduced into the recording chamber to recover for at least 2 hours. Two competing male flies with same age were introduced into the chamber simultaneously. To distinguish between genotypes, we marked the wings of flies in both the control and experimental groups one day before the behavioral assay. For example, in a setup with 50 pairs of flies, 25 pairs of control flies and 25 pairs of experimental flies were marked accordingly. The winner male was identified based on successful copulation with a female within a 30-minute observation period after the introduction of male competitors. All video recordings were manually analyzed to calculate the courtship parameters (courtship latency, courtship duration, and bending attempts).

The competitiveness of each male fly was assessed based on their chance of winning in the courtship competition. Briefly, each competing experiments carried out for single round was calculated as a single batch (typically one chamber group contains 32 pairs of competitors).

$$C=\frac{N_{experimental}-N_{control}}{N_{all}}$$

Where C is competitiveness, $N_{experimental}$ is the winning amount of experimental group, $N_{control}$

is the winning amount of control group, $N_{all}$ is the total pairs of this batch.

**Optogenetics**

Flies were collected within 24 hours after eclosion and single housed in food vials containing all-trans-retinal of 2.5mM final concentration. Flies were kept in dark and used for optogenetics experiments after fed with all-trans-retinal for 3 to 6 days.

For heart beat measurements, all flies received the corresponding light depending on the optogenetic tools (590nm for CsChrimson, 620nm for ReaChR) for 60 seconds.

For behavior activation, both experimental male and control male were introduced into the recording chamber under 10 s pulse light and 10s dark for 6 times (illustrated in Fig3, H) before the courtship assay.

**Female exposure**

The female exposure protocol was applied and modified from previous reported^3^. Briefly, the feminized male flies (donor flies) are obtained by expressing the sex determination gene *transformer* in the male oenocytes (Prom-E800-Gal4). 25 feminized male flies were then housed with 5 tester male flies in a food vial for 2 days (stress resistance assay) or until the death of all tester flies (life span assay).

**Stress Resistance and Life Span Assay**

In the stress resistance test, flies after co-housing with feminized males (see female exposure protocol above) were separated and transferred to new vials containing 1% agar for water supply. Dead flies were counted every 2~5 hours.

In the mating assay, tester flies were transferred into food vials either containing virgin female for copulation or food vials without female as control group. After 24 hours, tester flies from each group were transferred to new vials containing 1% agar. Recordings of death events were followed once tester flies were transferred into agar containing vials.

In the life span test, tester flies were doted by markers distinguishing from feminized male flies, and the dead tester flies were counted and removed each day. The number of feminized male flies was monitored and the dead ones were replaced by newly eclosed feminized flies to maintain a stable number of donor flies (25 for each vial).

**Calcium Imaging**

Male flies were raised 7-10 days before imaging. The brains were dissected in hemolymph as described before^4^ [108 mM NaCl, 5 mM KCl, 4 mM NaHCO_3_, 1 mM NaH_2_PO_4_, 15 mM ribose, and mM Hepes (pH 7.5); 300 mosM]. CaCl_2_ (2 mM) and MgCl_2_ (4 mM) were added to the AHL before use. Recordings were conducted by an Olympus BX51WI microscope with a 40X water immersion objective, an Andor Zyla camera, and a Uniblitz shutter. The fluorescent signals were recorded at 1 Hz and further analyzed using custom-made python scripts (supplementary information).

For functional connection assay, ATP was added to the saline with a final concentration of 2 mM after collecting stable signals from GCaMP expressing neurons.

The fluorescent change was calculated as follows: ΔF= (F_max_-F_0_)/ F_0_ where F_0_ was averaged for 50 frames before adding solution or chemicals.

**Immunohistochemistry**

Immunostaining was conducted according to previous protocol^4^. Brains and heart samples were dissected in dissection buffer [0.015% Triton X-100 in 1× phosphate-buffered saline (PBS)] and fixed in 4% para- formaldehyde (PFA) at room temperature on a nutator for 40 min. The tissues were then washed three times for 10 min in wash buffer (30% Triton X-100 in 1× PBS). Subsequently, the samples were blocked in block buffer (1× heat-inactivated normal goat serum with 30% Triton X-100 in 1× PBS) for 60 min at room temperature and incubated with primary antibody overnight at 4°C. On the second day, tissues were washed three times for 10 min in wash buffer and then incubated in secondary antibodies for 2-5 hours.

The following antibodies were used: rabbit anti-GFP (1: 250, Invitrogen), rabbit anti-RFP (1:250, Rockland) and mouse anti-nc82 (1:250, Developmental Studies Hybridoma Bank), Alexa 647-Phalloidin (1:2000, ATT Bioquest). Secondary antibodies were Alexa Fluor 555 goat anti-rabbit IgG (1: 250, Invitrogen), Alexa Fluor 488 goat anti-rabbit IgG (1: 250, Invitrogen), Alexa Fluor 647 goat anti-mouse IgG (1: 250, Invitrogen).

Images were acquired using a Nikon AX confocal microscope and Olympus IXplore spinning disk microscope.

**Mice**

**Mating preparation**

All female mice indicated as “estrus” in mating behaviors were induced to estrus period as previously described^5,6^. Briefly, female mice received 10μg of 17-β-estradiol benzoate (Sigma, cat#E8515) in 100 μl sesame oil on day -2, 5μg of 17-β-estradiol benzoate (Sigma, cat#E8515) in 50 μl sesame oil on day -1, and 50μg of progesterone (Sigma, cat# P0130) in 50 μl sesame oil on day 0. The behaviors are tested after 4-6 hours after day 0 injection.

Male mice were group housed initially and isolated 3-5 days before the behavior test.

**Female exposure process**

Female mice aged to 7-10 weeks after mating preparation were group housed and introduced randomly to males after habituation. All female mice were removed from the home cage of corresponding male mice after 1 hour.

**Vires injection**

Wild-type mice aged three to four weeks were injected with rAAV-cTNT::Tac1-RNAi (2 ×10^11^vg per mouse)or vehicle by intravenous injection. The sequence of Tac1 target has been previously reported^7^: GTTCTTTGGATTAATGGGCAA

A total volume of 100 µl virus was injected and mice were allowed to recover for 30 min before returning to home cage.

**Optical recording assay**

Heart rate was recorded with the MouseOx Pulse oximeter (s-collar clip, shaved neck; Starr Life Sciences) as preciously described^8^. Mice were shaved around the neck and acclimated to behaving with the collar sensor for at least five days during the standard handling procedure (Extend data figure 2, A). Hear rate was recorded during the whole female exposure process. Heart rate signals were recorded as beats per minute (bpm) at 15 Hz.

**ECG recording assay**

ECG recording of male mouse was done as previously described^9^. Briefly, male mice were anesthetized with Tribromoethanol (270 mg/kg) for surgery. Custom made electrodes were implanted to mouse with wire attached to chest muscle near heart (Extend data figure 2, B). Insulative part of the electrodes was anchored on the skull with glue (Krazy glue, Elmer’s Products Inc). Other part of the electrodes and wires exposed at the surface between PFA coat and electrodes were sealed by dental acrylic. The body temperature of mice during surgery was kept at 36 °C by a heating pad. After surgery all mice were allowed to recover for 5 days single housed in their home cage.

**Heart rate data analysis**

All heart rate data (including optical recording assay and ECG recording assay) were analyzed by custom made python scripts.

**Behavior test**

All male mice were maintained at home cage for mating behavior test and all test were performed in the dark cycle.

Male mice were introduced under camera for at least 10 min habituation before introduction of female mice. 100 ul of Saline or phenylephrine^10^ (PE, 1.5mg/ml) were injected to male mice before introduction of female mice. Following mating behaviors of male mice were recorded until ejaculation or 1 hour without visible ejaculation behavior after the entry of female mice.

Videos were manually annotated using software BORIS^11^, all behavior parameters including chasing, sniffing, mounting, intromission and ejaculation were annotated and calculated as courtship behaviors, behaviors from individuals are showed as representative results(Extend data figure 7, C).

**Reagent list**

| Strains | Source | Identifier |
| --- | --- | --- |
| w1118 | Lab stock | / |
| w[*]; ;red[1] e[1] htl[AB42]/TM3, P{ry[+t7.2]=ftz-lacC}SC1, ry[RK] Sb[1] Ser[1] | Bloomington Drosophila stock center (BDSC) | Stk#5370 |
| hand4.2-GAL4 | Xiushan Wu Lab, Hunan Normal University | / |
| w[*]; P{w[+mC]=Ugt36A1-GAL4.K}11C, P{w[+mC]=UAS-GFP.U}2 | BDSC | Stk#67608 |
| TER94-RNAi | THU stock | THU1058 |
| klf15-RNAi | THU stock |  |
| w[*] P{w[+mC]=EP}ppk23[G17320] | Yan Zhu Lab, Institute of Biophysics, Chinese Academy of Sciences | / |
| w[*]; ;ppk25-GAL4/TM6B | Yi Rao Lab, Peking University | / |
| w[-];UAS-kir2.1 | Chang liu Lab | / |
| w[-];UAS-TNT | Lab stock | M432 |
| UAS-tdTomato;CCAP-GAL4 | Lab stock | C227 |
| R71G01-LexA;CCAP-GAL4 | Lab stock | / |
| w[1118]; P{y[+t7.7] w[+mC]=GMR71G01-GAL4}attP2 | BDSC | Stk#39599 |
| w[*]; P{y[+t7.7] w[+mC]=UAS-ReaChR}attP40 | BDSC | Stk#53741 |
| w[*]; ;P{w[+mW.hs]=GawB}how[24B] | BDSC | Stk#1767 |
| y[1] v[1]; P{y[+t7.7] v[+t1.8]=TRiP.JF01338}attP2 | BDSC | Stk#31490 |
| w[1118]; P{w[+mC]=UAS-GFP.dsRNA.R}143 | BDSC | Stk#9331 |
| w[1118]; P{y[+t7.7] w[+mC]=GMR71G01-lexA}attP40 | BDSC | Stk#54733 |
| y[1] w[*]; Bl[1]/CyO, y[+]; P{w[+mC]=CCAP-GAL4.P}9 | BDSC | Stk#25686 |
| LexAoP-P2X2;UAS-GCaMP | YuFeng Pan Lab, Southeast University | / |
| CCHa1-RNAi | THU stock | THU03973.N |
| CCHa2-RNAi | THU stock | THU03595.N |
| ILP6-RNAi | THU stock | THU0985 |
| ILP7-RNAi | THU stock | THU1051 |
| ITP-RNAi | THU stock | THU2021 |
| Ms-RNAi | THU stock | THU2244 |
| nplp2-RNAi | THU stock | THU3239 |
| nplp3-RNAi | THU stock | THU3115 |
| nplp4-RNAi | THU stock | THU3143 |
| PPK-RNAi | BDSC | Stk#29571 |
| Piezo-RNAi | Lab stock | R202 |
| y[1] w[*];P{w[+mW.hs]=tin-Gal4.B}2 | BDSC | Stk#91538 |
| y[1] w[*];P{w[+m*]=tinC-Gal4.Delta4}12a | BDSC | Stk#92965 |
| TKR99D mutant | Yi Rao Lab CCT Line, Peking University | CY32-CG7887 |
| TKR99D-RNAi | THU stock | THU2675 |
| w[1118]; P{y[+t7.7] w[+mC]=20XUAS-IVS-CsChrimson.mVenus}attP2 | BDSC | Stk#55136 |
| UAS-GCaMP;R71G01-GAL4 | Lab stock | / |
| PromE(800)GAL4[4M],tub::GAL80ts | Yi Rao Lab, Peking University | / |
| UAS-traF | YuFeng Pan Lab, Southeast University | / |
| w[1118]; P{w[+mC]=Crz-GAL4.391}3M | BDSC | stk#51976 |
| Crz-R-RNAi | THU stock | THU2158 |
| UAS-ITPF | Wei Song Lab, Wuhan University | / |

**References**

1.	Mathis, A. *et al.* DeepLabCut: markerless pose estimation of user-defined body parts with deep learning. *Nat. Neurosci.* **21**, 1281–1289 (2018).

2.	Virtanen, P. *et al.* SciPy 1.0: fundamental algorithms for scientific computing in Python. *Nat. Methods* **17**, 261–272 (2020).

3.	Maures, T. J. *et al.* Drosophila Life Span and Physiology Are Modulated by Sexual Perception and Reward. *Science (80-. ).* **343**, 544–548 (2014).

4.	Zhang, L., Guo, X. & Zhang, W. Nutrients and pheromones promote insulin release to inhibit courtship drive. *Sci. Adv.* **8**, (2022).

5.	Inoue, S. *et al.* Periodic Remodeling in a Neural Circuit Governs Timing of Female Sexual Behavior. *Cell* **179**, 1393-1408.e16 (2019).

6.	Bayless, D. W. *et al.* Limbic Neurons Shape Sex Recognition and Social Behavior in Sexually Naive Males. *Cell* **176**, 1190-1205.e20 (2019).

7.	He, Z. X. *et al.* Nucleus Accumbens Tac1-Expressing Neurons Mediate Stress-Induced Anhedonia-like Behavior in Mice. *Cell Rep.* **33**, (2020).

8.	Klein, A. S., Dolensek, N., Weiand, C. & Gogolla, N. Fear balance is maintained by bodily feedback to the insular cortex in mice. *Science (80-. ).* **374**, 1010–1015 (2021).

9.	Hsueh, B. *et al.* Cardiogenic control of affective behavioural state. *Nature* **615**, 292–299 (2023).

10.	Yao, Y. *et al.* Cardiovascular baroreflex circuit moonlights in sleep control. *Neuron* **110**, 3986–3999 (2022).

11.	Friard, O. & Gamba, M. BORIS: a free, versatile open-source event-logging software for video/audio coding and live observations. *Methods Ecol. Evol.* **7**, 1325–1330 (2016).
